# IntelliProfiler: a research workflow for analyzing multiple animals with a high-resolution home-cage RFID system

**DOI:** 10.1101/2024.10.23.619967

**Authors:** Shohei Ochi, Hitoshi Inada, Noriko Osumi

## Abstract

Unbiased, scalable behavioral phenotyping that captures multi-animal interactions in home-cage settings is increasingly needed. Here we present “IntelliProfiler”, a research workflow consisting of data processing scripts that extract locomotor activity and pairwise proximity from a commercially available, previously validated, high-resolution radio frequency identification (RFID) floor plate. IntelliProfiler is not a standalone system; it operates on data acquired with the Phenovance floor plate and is not yet validated with other hardware configurations. The workflow reconstructs individual trajectories and positions of multiple mice, enabling long-term assessment of locomotor and social spacing. In proof-of-concept analyses, male mice placed in a novel cage environment maintained greater inter-animal distances than female mice, an effect that strengthened as group size increased. Aging reduced locomotor activity in a group size-dependent manner and altered proximity patterns. In addition, offspring of aged fathers (a paternal-aging autism spectrum disorder (ASD) model) exhibited hyperactivity and increased social distance relative to controls, consistent with ASD-related phenotypes and motivating further investigations. Together, these findings demonstrate that IntelliProfiler workflow provides a practical and versatile approach for screening group dynamics and quantifying complex social behaviors in neuroscience research.

## Introduction

Behavioral analyses using model animals are crucial for understanding neural functions and pathological states. Due to its ease of handling and suitability for genetic manipulations, the laboratory mouse is widely used as a model animal to study the neural bases of various conditions, including psychiatric, neurodevelopmental, and neurodegenerative disorders. Traditional methods for analyzing neural functions in mice involve assessing behaviors, such as locomotion, exploration, social interaction, and learning/memory, through specific tests, including the open field test, the elevated plus maze, the social interaction test, and the Morris water maze^1–4^. However, these traditional approaches come with several limitations.

A major limitation of traditional behavioral analyses is the low throughput of data acquisition, as most analyses are manually conducted over limited periods under artificial conditions. Typically, behavior is recorded for a single mouse in a specific experimental setup. For instance, the open field test measures general locomotor activity and anxiety-related behavior by placing a single mouse in an open arena and tracking its movements and time spent in different zones^1^. Similarly, the three-chamber social interaction test assesses sociability by comparing the time a mouse interacts with a novel mouse versus an empty chamber, offering insights into social preferences^5^. A more significant concern is that interactions between experimenters and animals often introduce biases that influence results^6–8^. Handling by experimenters can elevate heart rate, body temperature, spontaneous activity, and anxiety, potentially impacting memory and learning abilities^7,9^. Moreover, variations in behavioral data have been linked to the researcher’s gender and environmental conditions^8,10^. Therefore, there is a growing need for automated, unbiased systems to analyze behavior in semi-natural (home-caged) conditions to ensure more robust and reliable data.

Recent advances in behavioral analyses have increasingly focused on studying group dynamics in semi-natural environments. One widely adopted approach is the use of radio frequency identification (RFID) technology, which enables the tracking of multiple animals within a home-cage setting. Compared to manual video tracking, RFID systems offer continuous, high-throughput data acquisition with minimal human intervention. IntelliCage, a pioneering RFID-based system, utilizes RFID readers placed at the corners of a cage to automatically monitor the frequency and timing of animals entering chambers^11^. IntelliCage has been widely used to investigate spontaneous behavior and learning in group-housed mice^12,13^. Another RFID-based system, ECO-HAB, consists of four open fields connected by tubes, with RFID readers positioned at the junctions, allowing the study of free movement and social interactions among mice by tracking their movement between fields^14,15^. In addition, floor plate RFID systems have been developed to measure the locomotor activity of multiple animals simultaneously^16^. The first such system, “Trafficage”, was introduced in 2005 by NewBehavior and later commercialized by TSE Systems^17–19^, which was followed by “MultimouseMonitor”, being brought to market by PhenoSys^20^. More recently, a high-resolution RFID floor plate system was developed by Dr. Toshihiro Endo and his team and has been commercialized by Phenovance (Japan), enabling the precise tracking of multiple animals with improved temporal resolution^21^.

In this study, we developed IntelliProfiler, a research workflow comprising data processing scripts designed to analyze locomotor activity and social proximity using the commercially available, high-resolution RFID floor plate system described above.

While IntelliProfiler itself is purely a software-based workflow, here it was applied to data acquired from an RFID floor plate capable of tracking up to 16 mice with high spatial (5 x 5 cm grid) and temporal resolution^21^. This setup enables long-term, continuous monitoring of activity and proximity-based interactions with minimal researcher intervention. Using the IntelliProfiler workflow, we conducted 72-hour continuous behavioral analyses in large groups of male or female wild-type mice, comparing behavioral patterns between sexes across age groups to evaluate the effects of aging on social dynamics. We also examined a mouse model of autism spectrum disorder (ASD) derived from aged fathers^22,23^, assessing alterations of social behavior in offspring. Collectively, our findings demonstrate that IntelliProfiler workflow by provides a robust and efficient approach for investigating complex social behaviors and group dynamics in rodent models. Notably, IntelliProfiler does not include or control any RFID hardware; the number of animals and spatial resolution used in this study are determined entirely by the specifications of the external RFID board^21^.

## Results

### IntelliProfiler research workflow

The recording schema and physical configuration are shown in Fig. 1a and 1b. Raw data from the RFID floor plate system, including time, antenna ID, and transponder ID, were exported from the RFID floor plate (Supplementary Fig. 1a, c). These data were then converted into spatial coordinates and down-sampled to a per-second resolution using the IntelliProfiler workflow (Supplementary Fig. 1b, d). Subsequent calculations, statistical analyses, and figure visualization were performed on the converted data generated during the preprocessing step (Fig. 1c and Supplementary Fig. 2). Using this workflow, we conducted continuous 72-hour behavioral analyses on 8-week old male or female mice in groups of four, eight, and 16, recorded with RFID floor plate hardware (Fig. 1d). In the group of 16 male mice, one RFID tag was dislodged during the experiment; therefore data from 15 male mice were analyzed, following the approach described previously^24^.

**Fig. 1:**
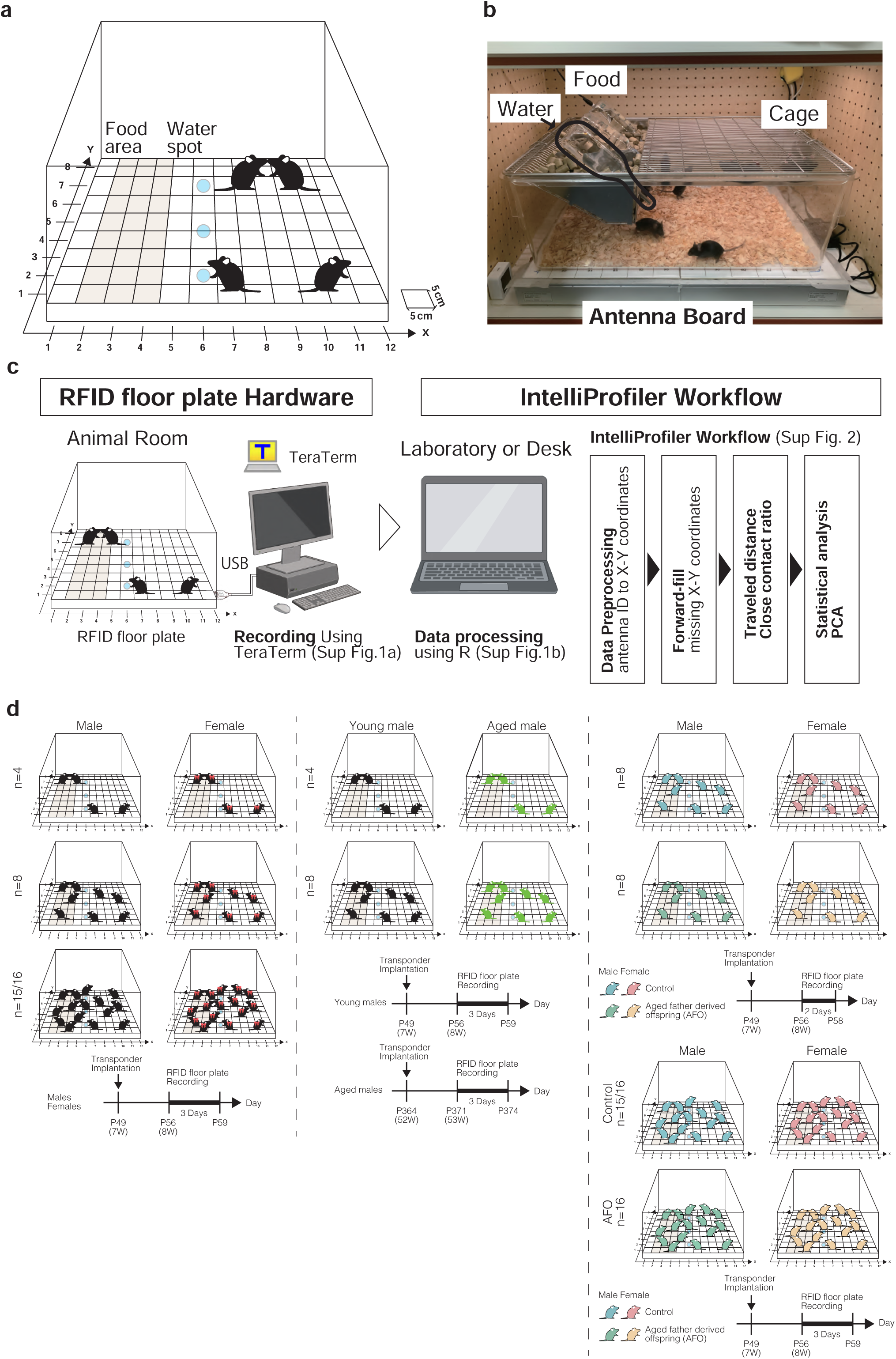
RFID floor plate automated behavioral measurement system and IntelliProfiler workflow data processing. **a** Schematic of RFID floor plate (Phenovance). The RFID floor plate system tracks the location of mice implanted with RFID tags via detectors placed in a 96-grid array at the base, with each grid measuring 5 cm x 5 cm. Location information is converted into XY coordinates (X=12, Y=8) using the IntelliProfiler workflow, based on RFID floor plate antenna IDs (1-96). **b** Photograph of the RFID floor plate. **c** Original data, capturing information on Time, Antenna ID, and Transponder ID from RFID floor plate hardware using TeraTerm. Raw hardware logging data were shown in Supplementary Fig. 1a. Processed data converted using the IntelliProfiler workflow. The processed data is refined to include Time, Antenna ID, Transponder ID, and XY coordinates. Decimal information is removed from the time data. If multiple individuals are detected on the same antenna ID simultaneously, their information is separated into different rows. **d** Behavior analysis was conducted using RFID floor plate over a continuous 72-hour period for male and female mice in groups of four-, eight- and 15/16. Young (8-week old) and aged (53-week old) male mice were analyzed in groups of four and eight. Aged father derived offspring (AFO) and control male and female mice were analyzed in groups of eight and 16.

### Locomotor activity

In the males, cumulative travel distances for every 2 hours exhibited the highest travel distance values in the four-mouse group than the eight- and 15-mouse groups during most of the time (Fig. 2a). The male mice exhibited the highest travel distance immediately after entry, dropping by less than half within 2 hours. Their travel distance increased again during the transition from light to dark period. In contrast, females showed less active and stable travel distances across almost all time points (Fig. 2b).

**Fig. 2:**
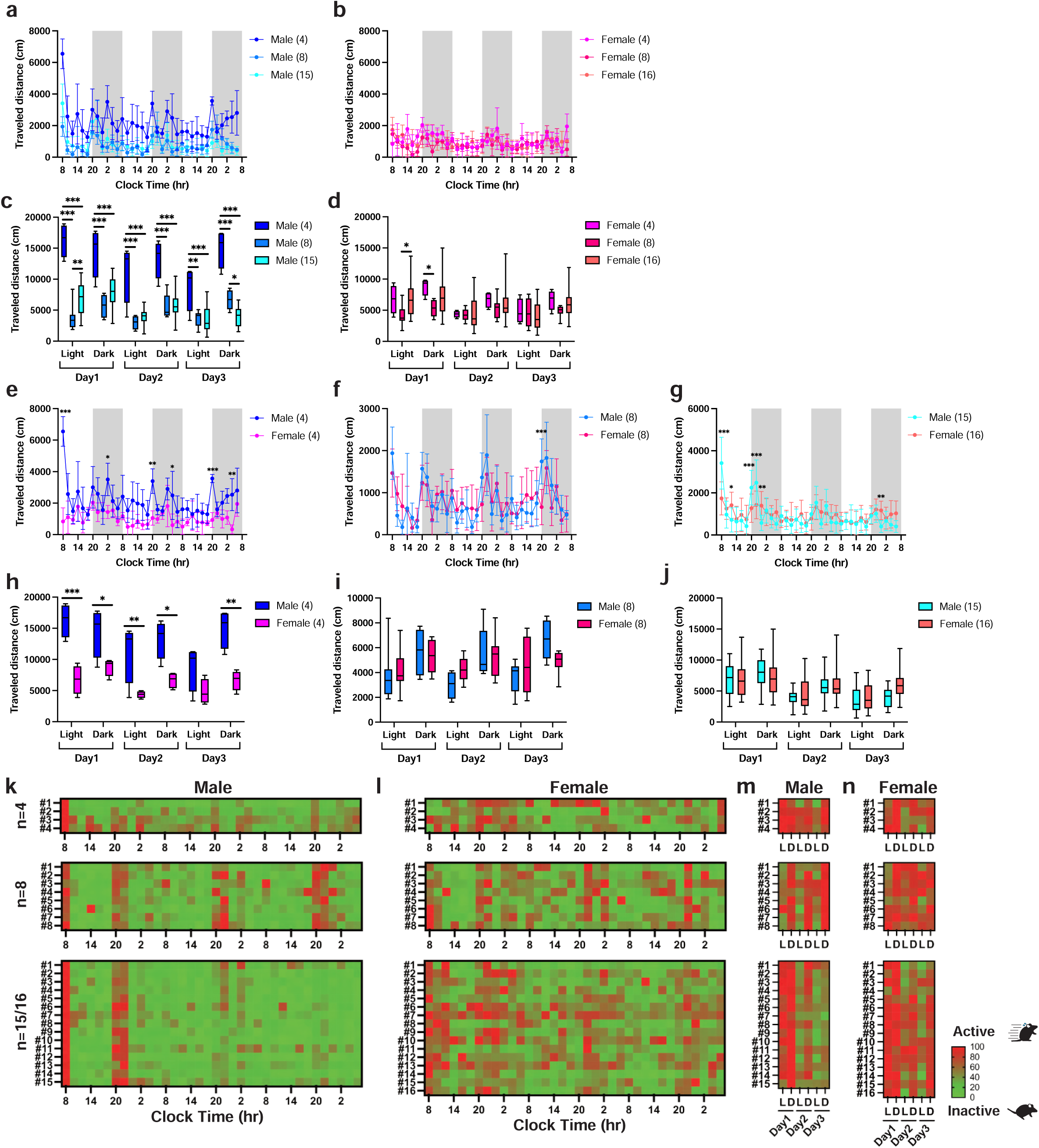
Activity analysis of group-housed mice using the IntelliProfiler workflow. **a-j** Cumulative travel distances measured in 2-hour intervals (**a, b, e-g**) and over 12-hour intervals (light-dark cycles) (**c, d, h-j**). **a-d** Analysis of locomotor activity based on group size, comparing groups of four, eight, and 15 males (**a, c**) and four, eight, and 16 female mice (**b, d**). **e-j** Analysis of sex differences in locomotor activity, comparing groups of four (**e, h**), eight (**f, i**) and 15-16 (**g, j**) male and female mice. **k-n** Heatmaps showing relative locomotor activity over 2-hour intervals (**k, l**) and 12-hour intervals (**m, n**) for groups of four, eight, and 15-16 mice. Values in the bar or line graphs are presented as mean ± SD, with box plots displaying min/max whiskers. Statistical analysis was conducted using Holm-Sidak method for multiple comparison. Significance levels are indicated as \**p* < 0.05, \*\**p* < 0.01, and \*\*\**p* < 0.001. *p* < 0.001 (**c**), *p* = 0.011 (**d**), *p* < 0.001 (**e**), *p* = 0.894 (**f**), *p* = 0.009 (**g**), *p* < 0.001 (**h**), *p* = 0.920 (**i**), *p* = 0.139 (**j**).

To understand the general trends of locomotor activity, we analyzed cumulative travel distances in 12-hour intervals. The males showed the highest travel distance just after their entry and significantly higher travel distance in the group of four mice than in the groups of eight and 15 mice (Fig. 2c). Small but significant differences were observed between the eight- and 15-mouse groups in the light period on day 1 and the dark period on day 3. On the other hand, females showed similar travel distances through the period among groups except for day 1 (Fig. 2d).

We then compared 2-hour interval activities to examine sex differences. In the four-mouse group, male mice showed higher travel distances throughout the period than females, with significantly higher travel distances observed at six time points in the dark periods (Fig. 2e). However, no periodic trend was observed in eight- and 15/16-mouse groups (Fig. 2f, g). In the12-hour intervals (Fig. 2h-j), males showed higher travel distances throughout the period in the four-mouse group than female mice except for the light period on day 3 (Fig. 2h). On the other hand, no noticeable sex difference was observed in the groups of eight and 15/16 mice (Fig. 2i, j).

The time course of relative locomotor activity for each mouse was visualized as a heatmap. Male mice showed a marked increase in activity immediately after the introduction and at the onset of the dark period (Fig. 2k). A pronounced contrast of locomotor activity was observed between light and dark periods over 3 days (Fig. 2m). In contrast, female mice displayed relatively more variable activity patterns over time (Fig. 2l), with some showing elevated activity even during the light period (Fig. 2n).

These findings suggest that male mice, particularly in smaller group sizes, exhibit higher locomotor activity, pointing to potential sex-specific behavioral responses to group size.

### Social interactions

To assess social interactions among group-housed mice, we tracked changes in the distance between two individual mice. The spatial relationships were classified into four categories based on proximities: “Same” - the other mouse occupied the same grid as the focal mouse; “Close” - the other mouse was in the adjacent grids; “Intermediate” - the other mouse was located beyond the surrounding grids; and “Away” (Fig. 3a). The transition in social distance ratios were recorded in 2-hour intervals for groups of four, eight, and 15 male mice and groups of four, eight, and 16 female mice.

**Fig. 3:**
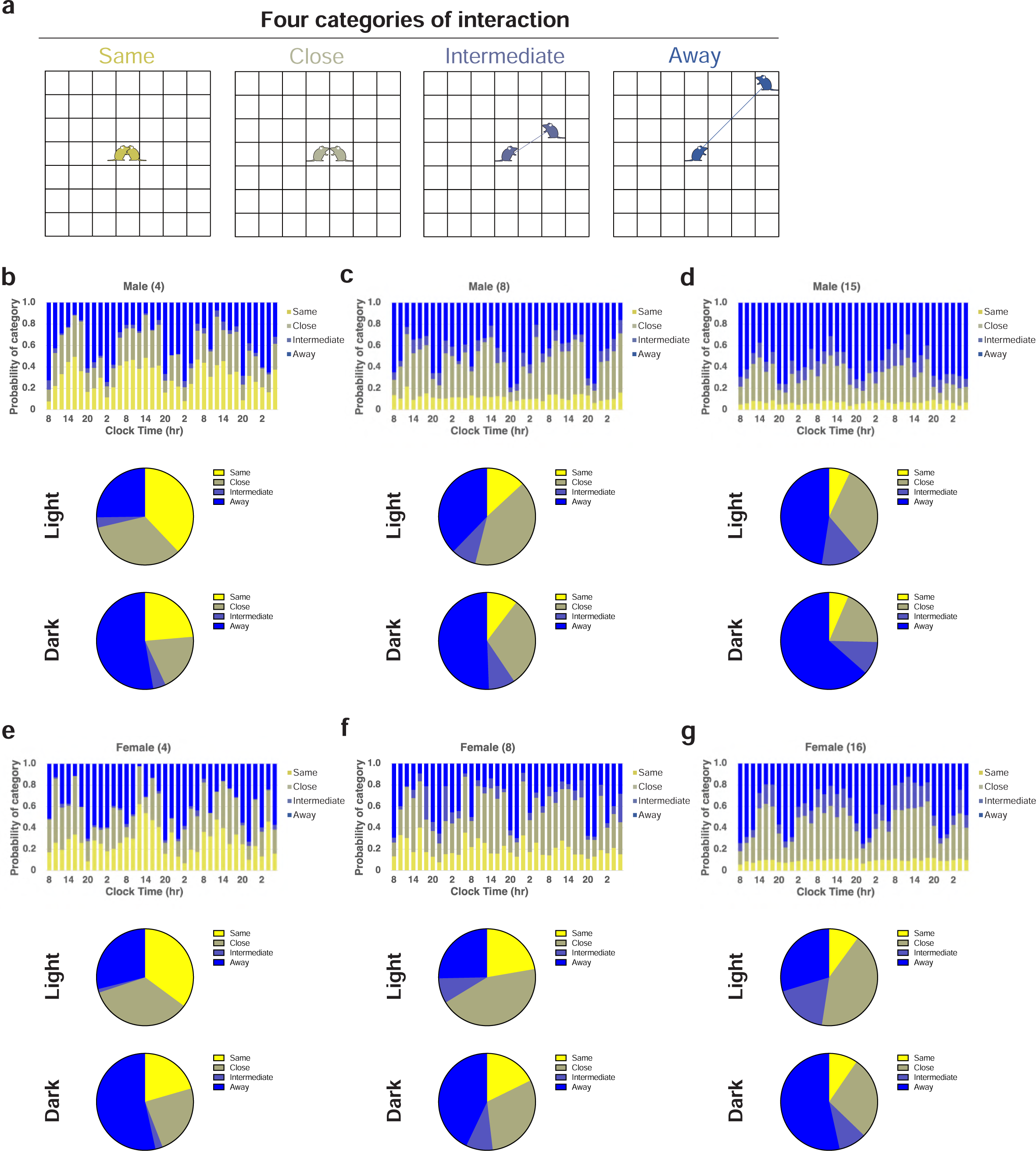
The spatial relationships among group-housed mice using the IntelliProfiler workflow. **a** Definition of the spatial relationships between two individual mice. When another mouse is in the same grid as the focal mouse (both depicted in yellow), the relationship is defined as “Same.” If the other mouse is in the surrounding grids (depicted in olive), it is defined as “Close.” If the other mouse is outside the surrounding grids (depicted in indigo), it is defined as “Intermediate.” If the other mouse is beyond the “Intermediate” zone (depicted in blue), the relationship is categorized as “Away.” **b-g** The 2-hour bin transition of proportions based on “Same,” “Close,” “Intermediate,” and “Away” for groups of four male mice (**b**), eight male mice (**c**), 15 male mice (**d**), four female mice (**e**), eight female mice (**f**), and 16 female mice (**g**). The average proportion of distance categories during the light period and the dark period is shown for each group. Chi-square *p*-values comparing the proportions between light and dark periods: *p* < 0.001 (**b**, four males), *p* = 0.256 (**c**, eight males), *p* = 0.114 (**d**, 15 males), *p* = 0.005 (**e**, four females), *p* = 0.049 (**f**, eight females), *p* = 0.007 (**g**, 16 females). The *p*-values of chi-square for the group size, *p* < 0.001 (**b-d**, males in the light phase), *p* = 0.002 (**b-d**, males in the dark phase), *p* < 0.001 (**e-g**, females in the light phase), *p* = 0.092 (**e-g**, females in the dark phase).

The proportion of “Same” increased during the light period and decreased during the dark period in the four-mouse group, but this pattern was not observed in the other groups (Fig. 3b-g). A similar trend was found for the “Close,” where its proportion rose in the light period and declined in the dark period across groups of four, eight, and 15 males and four, eight, and 16 females (Fig. 3b-g). The proportion of “Intermediate” was relatively low in the four-mouse group but increased with group size without showing a consistent periodic trend observed. In contrast, the proportion of “Away” decreased during the light period and increased in the dark period across all groups.

The average proportion of each social distance category during the light and dark periods was compared across groups. The proportion of “Same” decreased as group size increased. During the light period, the percentage of “Close” was higher than during the dark period across all groups, though there was no clear trend concerning group size.

The proportion of “Intermediate” increased with group size during the light period and was higher in the eight- and 15/16-mouse groups than the four-mouse group during the dark period, but similar between the eight-mouse and 15/16-mouse groups. In contrast, the proportion of “Away” was highest in the 15-male group during both periods, although no clear pattern emerged for the female groups. These analyses revealed qualitatively distinct patterns of social distance in the eight- and 15/16-mouse groups compared to the four-mouse group.

Significant differences in the proportions of each category between the light and dark periods were observed in the groups of four males and four, eight, and 16 females (Fig. 3b, e-g). However, no significant differences were found in the eight and 15 male groups (Fig. 3c, d). Regarding group size, significant differences were noted in the male groups during both the light and dark periods and in the female groups during the light period, but no significant differences were observed in the female groups during the dark period.

We then introduced the Close Contact Ratio (CCR), a metric to quantify social proximity to analyze social interactions. This ratio reflects the proportion of time two mice are either in the “Same” or “Close” (Fig. 4a). When CCR data were divided into 2- hour intervals for both male and female groups, males in the four-mouse group exhibited the highest CCR compared to the eight- and 15-mouse groups throughout the entire period. In male mice, the CCR was initially low immediately after entry, gradually increasing until 18:00, followed by a gradual decline until 20:00, with a similar pattern recurring from day 2 onward (Fig. 4b). In contrast, the females in the 16-mouse group showed a significantly lower CCR compared to the four- and eight-mouse groups (Fig. 4c), although no consistent time-dependent pattern was observed. We also analyzed the average CCR over 12-hour intervals. In male mice, the CCR decreased as the group size increased from four to eight to 15 mice during the light period (Fig. 4d). The 15-mouse group had the lowest CCR, while no significant difference was found between the four- and eight-mouse groups during the dark period. Similarly, female mice in the 16-mouse group tended to have the lowest CCR during the light period compared to the four- and eight-mouse groups, though no consistent differences were observed between the latter two groups (Fig. 4e). No clear trend was observed during the dark period in female mice.

**Fig. 4:**
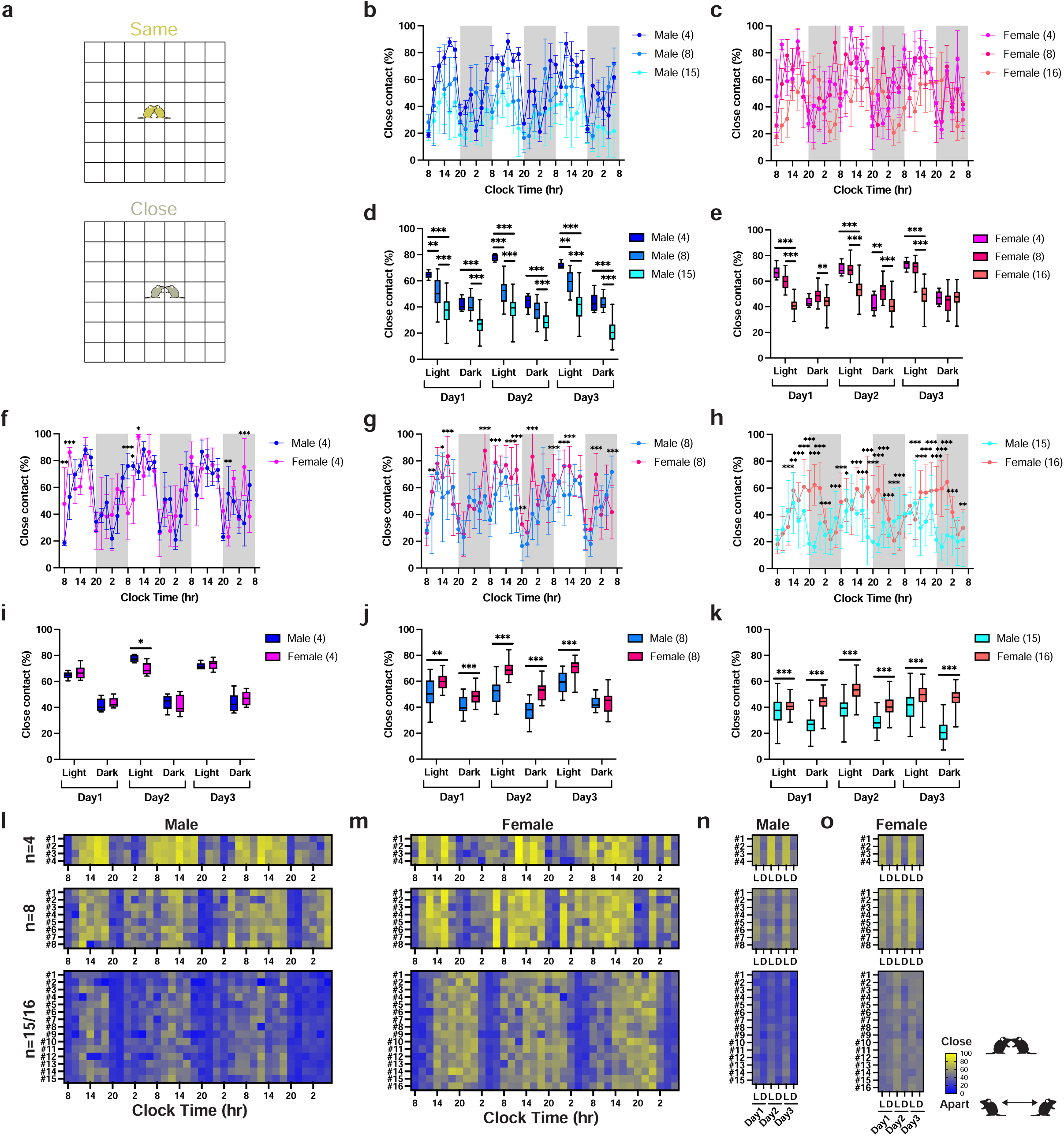
Close contact between two individuals among group-housed mice analyzed using the IntelliProfiler workflow. **a** Definition of close contact, where two mice either in the “Same” grid (depicted in yellow) or in adjacent grids, classified as “Close” (depicted in olive). **b-k** Average Close Contact Ratio (CCR) measured in in 2-hour intervals (**b, c, f-h**) and 12-hour intervals (**d, e, i-k**). **b-e** Analysis of CCR based on group size, comparing groups of four, eight, and 15 male mice (**b, d**) and four, eight, and 16 female mice (**c, e**). **f-k** Analysis of sex differences in CCR, comparing groups of four (**f, i**), eight (**g, j**) and 15/16 (**h, k**) male and female mice. **l-o** Heatmaps showing the time course of CCR over 2-hour intervals (**l, m**) and 12-hour intervals (**n, o**) for groups of four, eight, and 15-16 mice. Values in the bar or line graphs are presented as mean ± SD, with box plots showing min/max whiskers. Statistical analysis was conducted using Holm-Sidak method for multiple comparisons. Significance levels are indicated as \**p* < 0.05, \*\**p* < 0.01, and \*\*\**p* < 0.001. *p* < 0.001 (**b**), *p* < 0.001 (**c**), *p* = 0.882 (**f**), *p* < 0.001 (**g**), *p* < 0.001 (**h**), *p* = 0.873 (**i**), *p* < 0.001 (**j**), *p* < 0.001 (**k**).

We also examined the sex differences in CCR using 2-hour intervals in four-, eight-, and 15/16-mouse groups. In the four-mouse group, females showed a significantly higher CCR during the initial 4 hours (Fig. 4f). However, CCR fluctuated between males and females throughout the remaining time points, suggesting that consistent sex differences in social proximity might not be present in this group. In contrast, in the eight- and 15/16-mouse groups, females consistently exhibited higher CCR than males at most time points, indicating more pronounced sex differences in social proximity (Fig. 4g, h). To assess the overall trend, we analyzed the CCR in 12-hour intervals. In the four-mouse group, no clear sex differences in CCR were observed, except during the light period on day 2, when females had a higher CCR (Fig. 4i). However, in the eight- mouse group, females had a higher CCR than males across most periods, except during the dark period on day 3 (Fig. 4j). Similarly, in the 16-mouse group, females consistently exhibited a higher CCR than males across all the periods (Fig. 4k).

The time course of CCR, visualized as a heatmap for individual mice, revealed a strong periodic trend in the males, with CCRs peaking during the light period and decreasing during the dark period (Fig. 4l). In contrast, the female groups generally displayed higher CCRs for longer durations compared to males (Fig. 4l, m). The periodic patterns of CCRs were less pronounced in the 15/16 mouse groups, though the trend remained more apparent in males than females (Fig. 4l-o). These findings suggest that sex-specific social dynamics become more prominent in larger groups.

### Aging effect

Since aging has been reported to affect many behavioral aspects, including locomotor activity and social interaction^25,26^, we compared young (8-week-old) and aged (53-week-old) male mice in groups of four and eight (Fig. 1d and Supplementary Fig. 3-5). Aged males in the four-mouse group showed reduced social proximity, while those in the eight-mouse group exhibited prolonged movement durations, indicating that aging impacts activity and social behavior, particularly with group size.

### Application into ASD model mice

We further applied IntelliProfiler workflow to analyze behavior in a mouse model of ASD induced by advanced parental aging, using 8-week-old male and female offspring from aged (AFO) and control fathers^22,23^, housed in groups of eight or 16 (Fig. 1d).

In both group sizes, AFO males showed consistently increased travel distances across nearly all time points, while AFO females showed elevated activity at two-thirds of the time points (Fig. 5a-f). These effects were more pronounced in the 16-mouse group, with AFO males exhibiting greater cumulative distances, especially on Day 2 and during the light phase (Fig. 5g-l). Heatmaps indicate peak activity after cage introduction and at the onset of the dark phase, with AFO males displaying prolonged activity over time (Fig. 5m–t), suggesting altered temporal patterns.

**Fig. 5:**
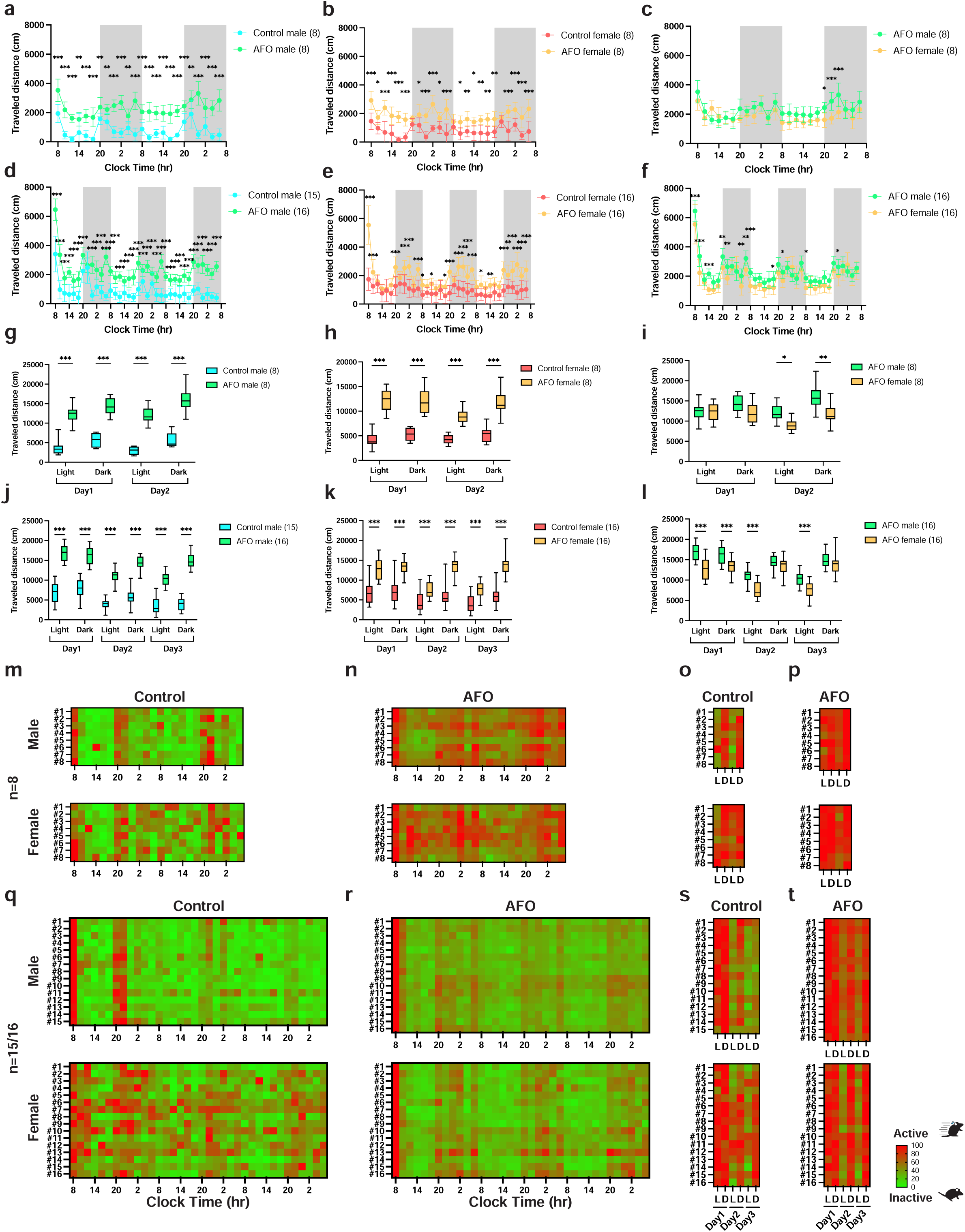
Activity analysis of group-housed male and female aged father derived offspring (AFO) mice using the IntelliProfiler workflow. **a-f** Cumulative travel distances measured in 2-hour intervals for 2 days (**a-c**) and for 3 days (**d-f**) and over 12-hour intervals for 2 days (**g-i**) and for 3 days (**j-l**). **a-l** Analysis of locomotor activity based on group size, comparing groups of eight and 15/16 aged father derived offspring (AFO) and control mice. **a-l** Analysis of parental aging effects in locomotor activity, comparing groups of eight and 15/16 male AFO and control mice (**a, d, g, j**), groups of eight and 16 female AFO and control mice (**b, e, h, k**), groups of eight and 16 male and female AFO mice (**c, f, i, l**). **m-t** Heatmaps showing relative locomotor activity over 2-hour intervals (**m, n, q, r**) and 12-hour intervals (**o, p, s, t**) for groups of eight and 16 mice. Values in the bar or line graphs are presented as mean ± SD, with box plots displaying min/max whiskers. Statistical analysis was conducted using Holm-Sidak method for multiple comparisons. Significance levels are indicated as \**p* < 0.05, \*\**p* < 0.01, and \*\*\**p* < 0.001.

Spatial proximity analysis revealed that AFO mice spent less time in close proximity and more time in intermediate or distant positions compared to control mice (Fig. 6a–i). CCR values were lower in AFO mice, especially in males during the dark phase and in females across both phases, with effects amplified in larger groups (Fig. 7a–g). AFO females showed higher CCRs values than males during the light phase (Fig. 7d, g, m), while cumulative CCRs confirmed reduced social proximity in AFO mice (Fig. 7h–l). CCR heatmaps further revealed greater individual variability within the AFO groups (Fig. 7n–u).

**Fig. 6:**
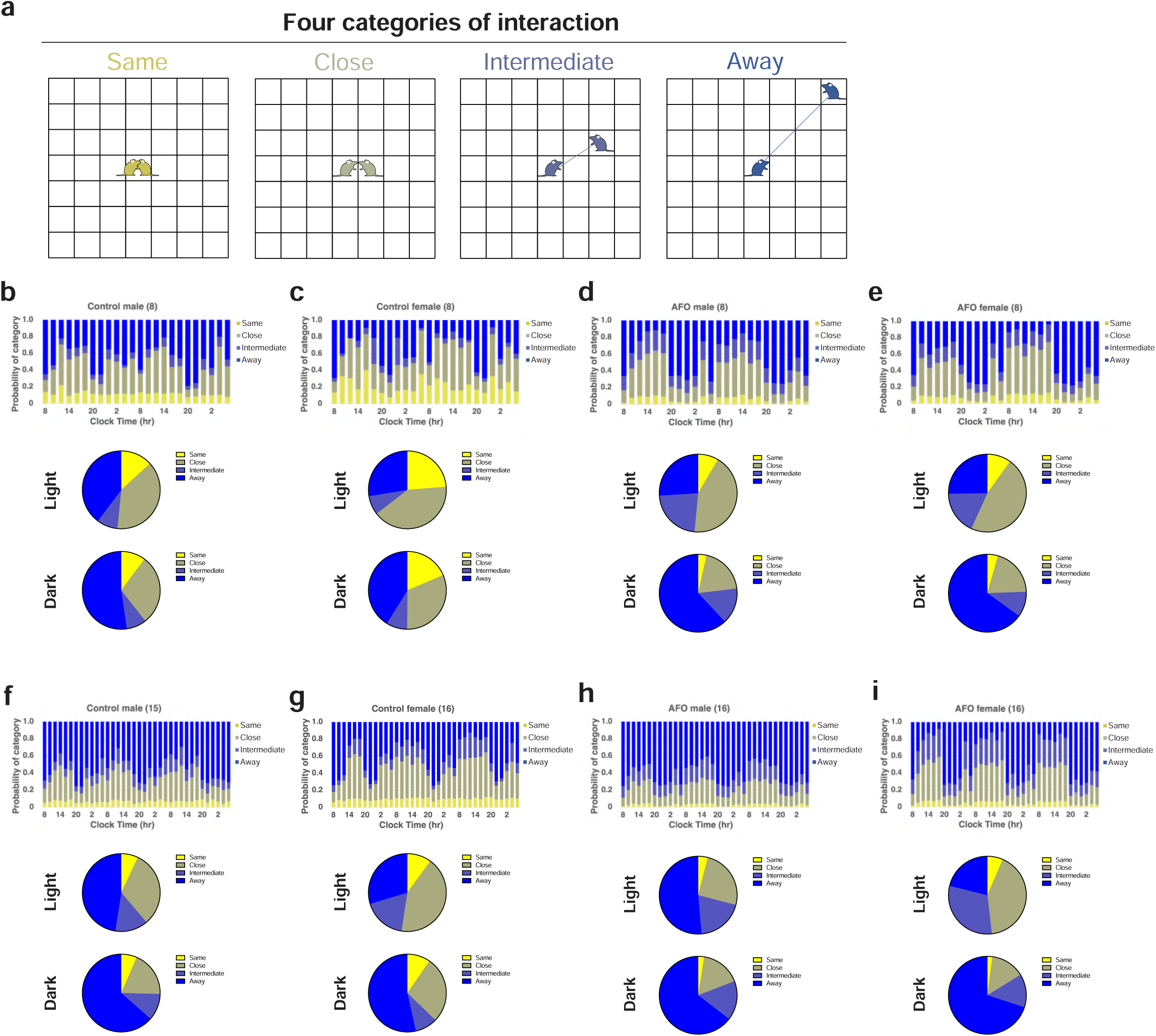
The spatial relationships among group-housed male and female AFO mice using the IntelliProfiler workflow. **a** Definition of the spatial relationships between two individual mice. When another mouse is in the same grid as the focal mouse (both depicted in yellow), the relationship is defined as “Same.” If the other mouse is in the surrounding grids (depicted in olive), it is defined as “Close.” If the other mouse is outside the surrounding grids (depicted in indigo), it is defined as “Intermediate.” If the other mouse is beyond the “Intermediate” zone (depicted in blue), the relationship is categorized as “Away.” **b-e** The 2-hour bin transition of proportions based on “Same,” “Close,” “Intermediate,” and “Away” for groups of eight male control mice (**b**), eight female control mice (**c**), eight male AFO mice (**d**), eight female AFO mice (**e**), 15 male control mice (**f**), 16 female control mice (**g**), 16 male AFO mice (**h**), 16 female AFO mice (**i**). **b-i** The average proportion of distance categories during the light period and the dark period is shown for each group. Chi-square *p*-values comparing the proportions between light and dark periods, *p* = 0.359 (**b**, eight male control mice), *p* = 0.215 (**c**, eight female control mice), *p* < 0.001 (**d**, eight male AFO), *p* < 0.001 (**e**, eight female AFO), *p* = 0.114 (**f**, 15 male control mice), *p* = 0.007 (**g**, 16 female control mice), *p* = 0.050 (**h**, 16 male AFO), *p* < 0.001 (**i**, 16 female AFO).

**Fig. 7:**
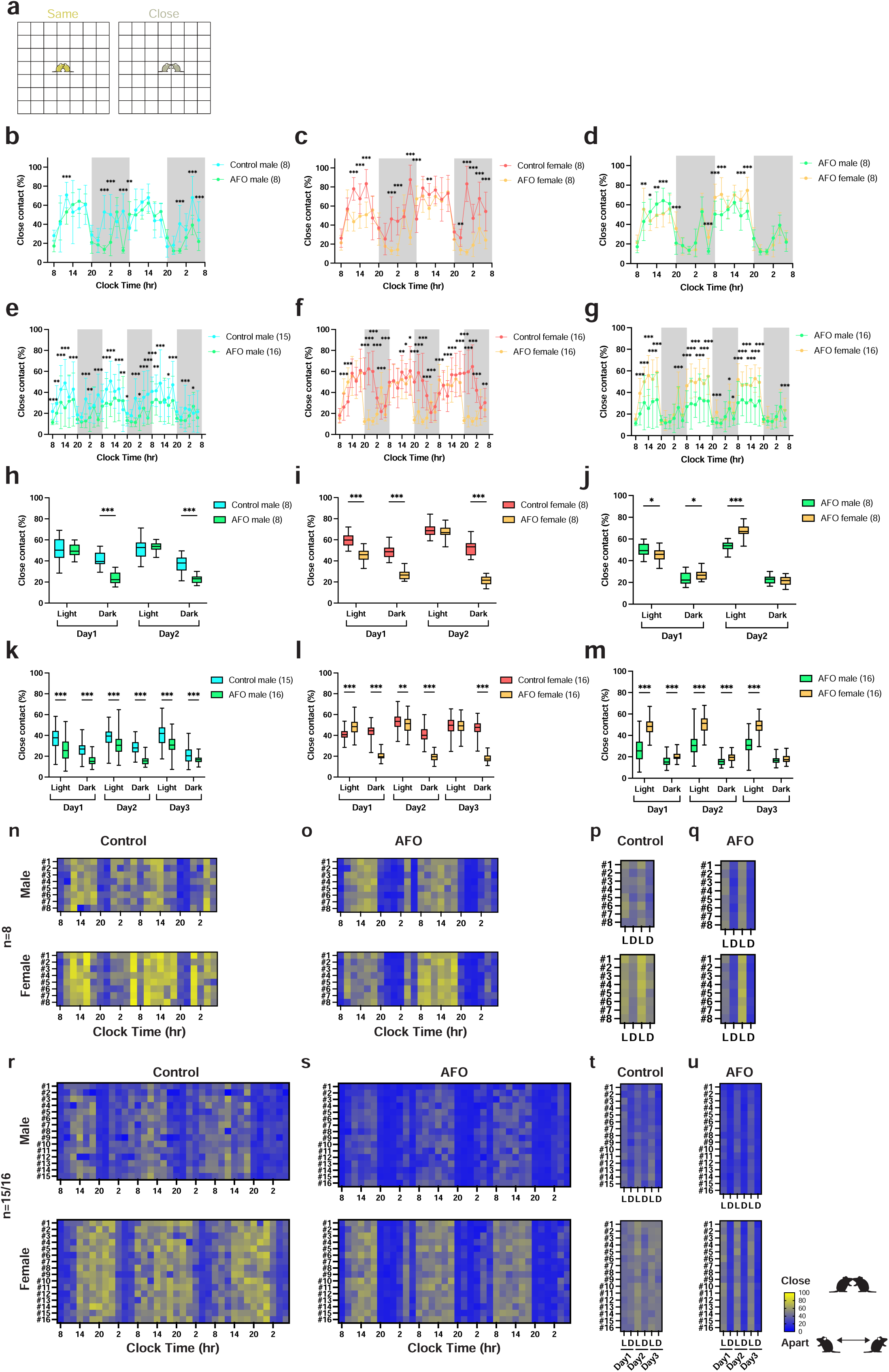
Close contact between two individuals among group-housed male and female AFO mice analyzed using the IntelliProfiler workflow. **a** Definition of close contact, where two mice either in either “Same” (depicted in yellow) or in adjacent grids, classified as “Close” (depicted in olive). **b-h** Average Close Contact Ratio (CCR) measured in 2-hour intervals (**b-g**) and 12-hour intervals (**h- m**). **b, e, h, k** Analysis of CCR based on group size, comparing groups of eight and 15/16 male AFO and control mice. **c, f, i, l** Analysis of CCR based on group size, comparing groups of eight and 16 female AFO and control mice. **d, g, j, m** Analysis of CCR based on group size, comparing groups of eight and 16 male AFO and female AFO. **n-u** Heatmaps showing the time course of CCR over 2-hour intervals (**n, o, r, s**) and 12-hour intervals (**p, q, t, u**) for groups of eight and 15/16 mice. Values in the bar or line graphs are presented as mean ± SD, with box plot showing min/max whiskers. Statistical analysis was conducted using Holm-Sidak method for multiple comparisons. Significance levels are indicated as \**p* < 0.05, \*\**p* < 0.01, and \*\*\**p* < 0.001. *p* < 0.001 (**c**), *p* = 0.338 (**d**), *p* = 0.072 (**e**), *p* < 0.001 (**f**), *p* = 0.007 (**g**), *p* = 0.109 (**h**).

### Characterization of group behaviors

To investigate the behavioral characteristics of each group, we conducted principal component analysis (PCA) using four key parameters. First, we defined normalized activity as the activity at time *t*, divided by the maximum activity value recorded over Days 1-3, and multiplied by 100 to standardize the data. This allowed us to calculate the normalized activity in the light (AL) and dark (AD) periods by averaging activity values at each time point over three days. We selected four parameters for PCA: sociability, measured as CCR values during the light (SL) and dark (SD) periods (parameters 1 and 2, respectively), and normalized activity during the light (AL) and dark (AD) periods (parameters 3 and 4, respectively). The PCA results with estimated clusters are displayed in Fig. 8. Correlations among these parameters are shown in Supplementary Fig. 6-8. The number of clusters was determined using the Elbow method^27^ and further assessed through Silhouette plots^28^ (Supplementary Fig. 9-11).

**Fig. 8:**
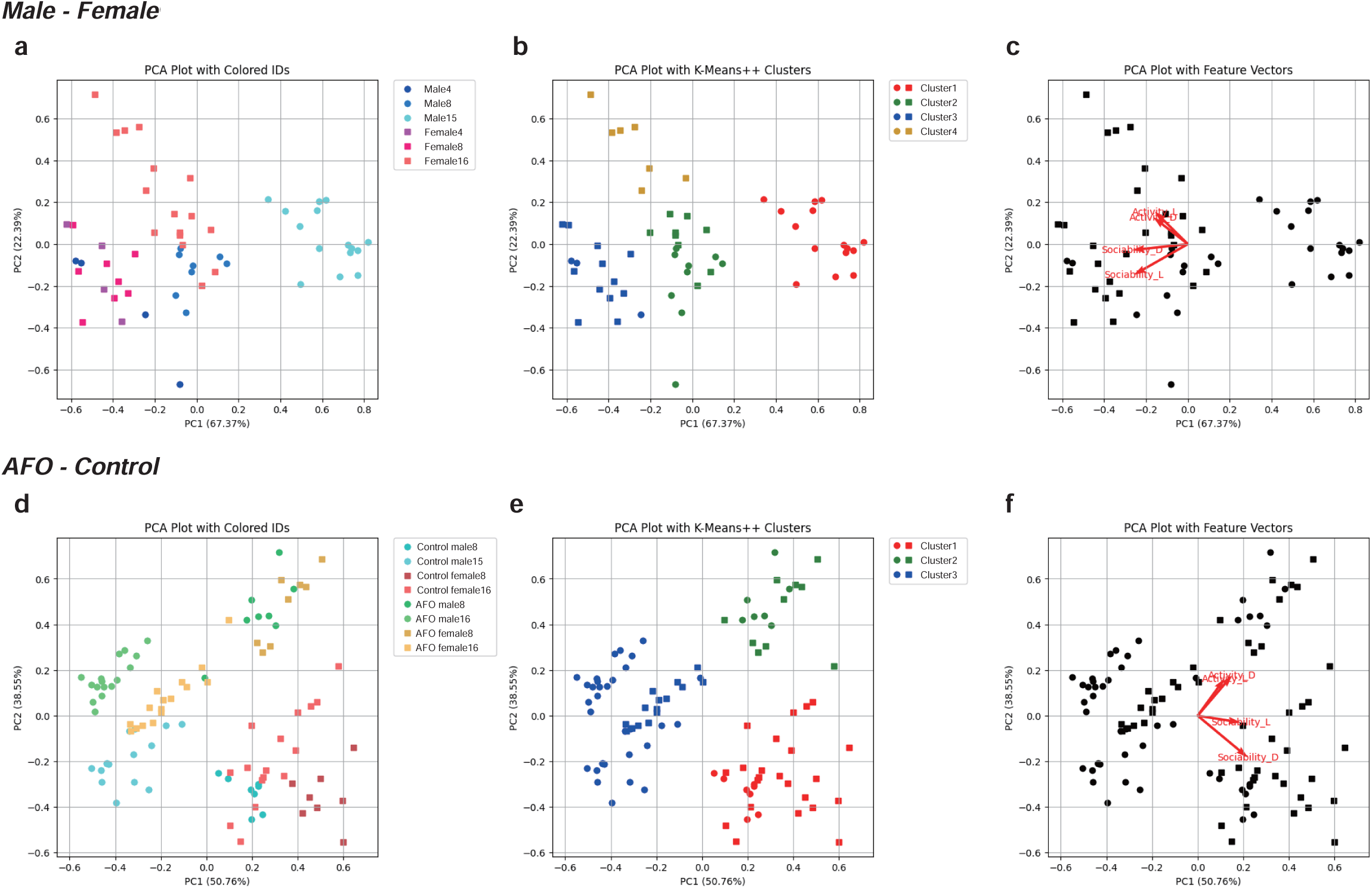
Principal component analysis of group behavior using the IntelliProfiler workflow. **a-f** Principal component analysis (PCA) of group behavior in four, eight, and 15/16 male and female mice (**a-c**) and in AFO and control mice in groups of eight and 15/16 (**d-f**). **b, e** K-means clustering in PCA plots. **c, f** Analysis of feature vectors, focusing on Close Contact Ratio (CCR) and relative activity during the light and dark periods.

The PCA revealed that the male mice in the 15-mouse group formed a well-defined separated cluster whereas the male and female mice in the four-mouse groups and female mice in the eight-mouse group clustered together (Fig. 8a). Female mice in the 16-mouse group split into two clusters, with one cluster aligning with the male mice in the eight-mouse group (Fig. 8b). In the feature space, AL and AD vectors pointed in the same direction, indicating similar patterns of activity, while SL and SD vectors diverged, reflecting different sociability dynamics between light and dark periods (Fig. 8c). In terms of aging effects, young and aged mice in the four-mouse groups were largely distributed, whereas those in the eight-mouse groups clustered near the origin (0,0) or in the negative X direction (Supplementary Fig. 10a). K-means clustering separated the four- and eight-mouse groups, indicating that aging influenced both activity and sociability in distinct ways (Supplementary Fig. 10b, c).

In the PCA analysis comparing AFO and control mice (Fig. 8d–f), AFO mice were primarily distributed along the negative axis of PC1, while control mice shifted toward the positive side. Control females predominantly clustered in cluster 1, whereas AFO females were classified into clusters 2 or 3, indicating a clear group separation (Fig. 8e). In the feature space, AL and AD vectors aligned, while SL and SD diverged (Fig. 8f), highlighting distinct contributions of locomotor and social features.

### Network analysis of social interactions

We explored how group size influences social interaction networks (Fig. 9). Qualitatively, there was no noticeable difference in network structure between male and female mice in the four- and eight-mouse groups (Fig. 9). However, in the 15/16-mouse groups, male mice exhibited increased social proximity and developed more expanded networks, influenced by light-dark circadian rhythms. In contrast, female mice maintained relatively uniform network structures (Fig. 9). When comparing young and aged male mice in the four-mouse groups, no significant differences in network structure were observed (Supplementary Fig. 12). In the eight-mouse groups, however, young male mice displayed increased social proximity and formed diverse networks that fluctuated with light-dark cycles, while aged male mice maintained more stable, uniform networks with only slight variation corresponding to circadian rhythms (Supplementary Fig. 12). When comparing AFO and control mice in both the eight- and 16-mouse groups, the social network structures of AFO males and females appeared more expanded than those of their control counterparts, with the more pronounced alterations observed in AFO female groups during the dark phase (Fig. 10). These findings suggest that male and female mice create distinct social environments and that aging may impact the complexity and heterogeneity of the networks in larger groups.

**Fig. 9:**
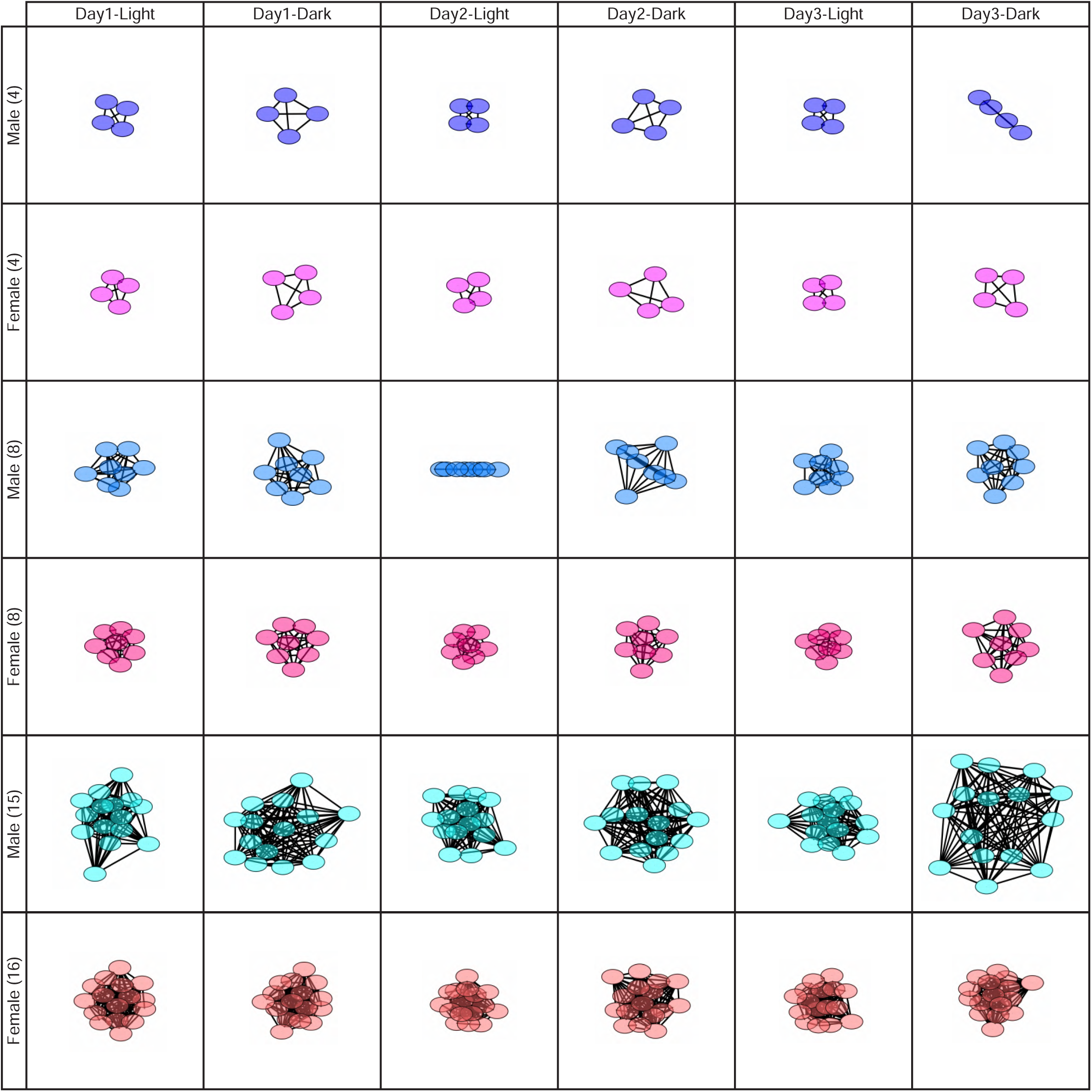
**Network analysis of social interaction in the male and female mice using the IntelliProfiler workflow.** Network analysis of social interactions during the light period on day 1 and the dark period on day 3 in groups of four, eight, and 15/16 male and female mice. Nodes represents individual mice, and edges reflects the relative social contacts between two individuals. The network graph was generated using Cytoscape. **Supplementary** Fig. 12 provides additional network diagrams, where unique color code for each individual highlighted the individual-specific social dynamics within the network.

**Fig. 10:**
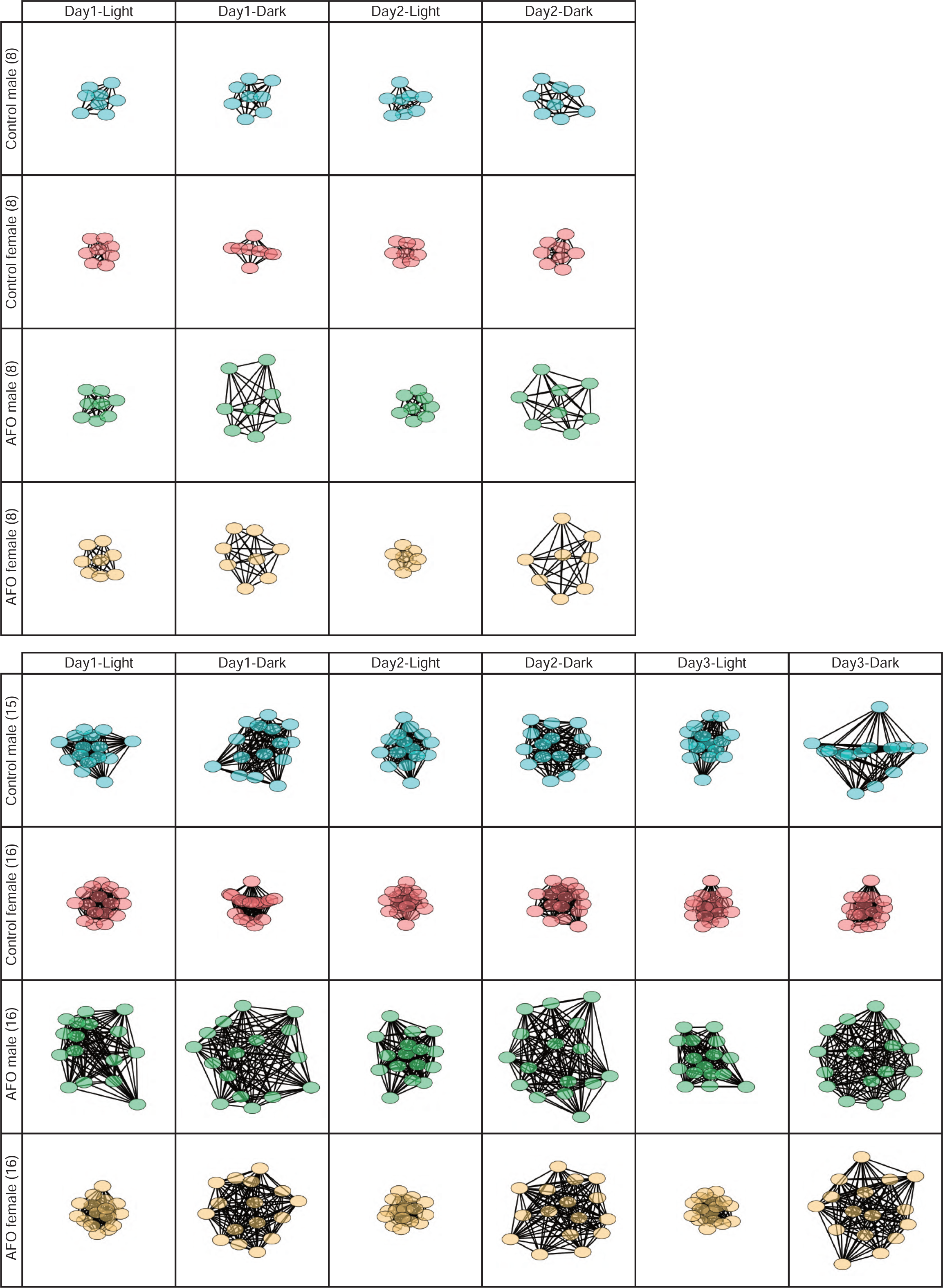
**Network analysis of group interaction in male and female AFO mice using the IntelliProfiler workflow.** Network analysis of social interactions during the light period on day 1 and the dark period on day 3 in groups of four and eight and 15/16 male and female AFO and control mice. Each circle represents an individual mouse, and the proximity of the lines indicates the relative social distance between two individuals. The network graph was generated using Cytoscape.

### Analysis of individual social interactions

Lastly, we focused on individual differences in social interactions through a pairwise analysis of CCR. No obvious individual differences were observed between sexes in the four- and eight-mouse groups (Supplementary Fig. 13-15). However, in the 15/16-mouse groups, two males (mouse ID #1 and #2) exhibited lower CCR scores than the others. Regarding the aging effect, aged males showed lower CCR scores and more ambiguous, less defined time-dependent patterns than young males (Supplementary Fig. 16-18). These findings suggest that individuality may develop progressively with aging in males, potentially contributing to the observed social dynamics.

## Discussion

In this study, we implemented “IntelliProfiler,” a research workflow comprising data processing scripts, which were developed to process raw positional data collected via a commercially available RFID floor plate system^21^ in combination with RFID tag implantation. Unlike fully integrated platforms such as “IntelliR”^29^, the current IntelliProfiler workflow is semi-automated and requires users to have basic proficiency in R and other analysis tools. Although not yet packaged as a standalone application, the workflow offers flexibility and transparency, making it well suited for research adaptation. IntelliProfiler itself contains no hardware components and is compatible with position data from high-resolution RFID boards, such as the one developed by Phenovance. The antenna board used in our study has previously been described as “a high-resolution RFID floor plate” in a recent review of the IntelliCage system^21^.

By applying IntelliProfiler workflow, we obtained novel insights into how group size, sex, and aging influence on locomotor activity and social proximity, thereby providing a more comprehensive understanding of semi-natural behaviors in group-housed mice.

Although RFID tag implantation is an invasive procedure and may have minor post-surgical effects, it is already used for IntelliCage and other systems. Our protocol enables rapid and well-tolerated tagging, supporting longitudinal behavioral studies under home-cage conditions with minimal disruption.

One of our key findings was the impact of group size on behavior. Using the IntelliProfiler workflow, we detected significant differences in both activity levels and social proximity across group sizes. Traditional behavior paradigms, which are typically designed for individual animals^30,31^, may overlook the complexities that emerge in group settings. Recent work has underscored the importance of studying animals under ’natural conditions’^32^. By enabling continuous monitoring of authentic locomotor activity in semi-natural conditions, the IntelliProfiler workflow provides a more comprehensive view of how group size shapes both individual activity patterns and social dynamics.

We introduced CCR as a novel social parameter to measure social proximity, representing a significant advancement in behavioral analysis (Fig. 4). Traditional social interaction tests offer limited insight into short-term interactions, but the CCR provides a more detailed, long-term view of group dynamics. We found that females tended to form stable clusters, while males exhibited more fluid, group size-dependent social patterns (Fig. 4, 8, 9).

While CCR serves as a useful proxy for spatial proximity and potential sociability, it is important to recognize that such proximity can also result from non-social factors. High CCR value may reflect a range of behavioral motivations, including social affiliation, thermoregulation, or reduced locomotor activity due to aging or pathology. Although our study maintained consistent environmental conditions (e.g., temperature, cage size, light/dark cycles), the interpretation of CCR as a social metric should be made with caution and in context. Future studies incorporating temperature manipulation or single-housing controls will help disentangle these contributing factors.

Previous studies using depth-sensing or RFID-assisted video tracking systems have enabled detailed assessment of social behaviors such as approaching, avoiding, or attacking by detecting animal orientation with the high spatial resolution^33–36^. However, these systems are typically limited in the number of animals that can be tracked simultaneously. In contrast, the CCR metric used in our study reflects spatial proximity but does not directly capture specific social actions. Still, time-resolved CCR trajectories may support broader analyses of such social interactions. Notably, CCR analysis avoids identity-swapping errors that can occur in video tracking with large groups.

Aging affected both locomotor activity and social interactions, with aged males showing more variable and less distinct CCR patterns compared to young males (Supplementary Fig. 4-5). In the eight-mouse group, aged males exhibited higher activity levels than younger males (Supplementary Fig. 3), contrasting with studies using open field test that reported reduced or similar travel distances in aged mice^26,37^. This discrepancy likely stems from differences in the experimental context—short-term, single-animal testing in a novel environment versus long-term monitoring in group-housed, semi-natural settings. In terms of social behavior, our findings align with previous reports of reduced contact frequency and shorter interaction durations in aged mice^26^.

To assess behavioral variability, we analyzed coefficient of variation (CV) for both travel distance and CCR across experimental conditions (Supplementary Fig. 21-24). CVs for travel distance increased with group size in both sexes (Supplementary Fig. 21e–f), while aged males showed similar variability to young males in smaller groups (Supplementary Fig. 21g–l), suggesting that aging had limited impact on variability. A similar trend was observed for CCR: male mice showed greater variability with increasing group sizes, whereas females maintained more consistent (Supplementary Fig. 23e–f). AFO mice showed lower variability in travel distance compared to control mice (Supplementary Fig. 22), although both groups exhibited increased CCR variability with larger group sizes (Supplementary Fig. 24). These results suggest that behavioral variability is shaped by group size and sex, and epigenetic factors such as advanced paternal age. Elevated CVs in large male groups may reflect greater heterogeneity or emerging social stratification. A limitation of this study is that the AFO and control mice were generated, housed and recorded in different facilities, which may have introduced confounding effects.

Our findings show that AFO mice, a model of ASD, exhibited increased locomotor activity and reduced social proximity compared to control mice, particularly in larger groups (Fig. 5–7). While previous studies reported no difference^38^ or reduced activity in AFO mice^39^, our group-housed analysis revealed increased locomotor activity, suggesting a context-dependent phenotype that emerges only under ecologically valid conditions. Regarding social behavior, prior studies have shown reduced sociability in AFO mice, especially with advanced paternal age^38–40^. Consistent with these reports, IntelliProfiler workflow detected lower CCR values in AFO mice, particularly in larger groups—a pattern not observed in our previous study^41^. These results underscore the utility of IntelliProfiler workflow as a context-sensitive platform for detecting subtle behavioral phenotypes in neurodevelopmental disorder models.

Our study was not intended to provide a comprehensive analysis of sex differences per se, but rather highlights the interaction between age, genotype, and social context in shaping group dynamics. While some sex-related effects have been previously described^42^, our results extend these findings by revealing distinct behavioral trajectories in aged males and AFO mice under RFID floor plate conditions, incorporating both physical activity and social proximity metrics. These observations serve as illustrative examples of how the IntelliProfiler workflow can capture complex group dynamics, including those influenced by age and sex. Although some patterns were consistent with prior literatures, the workflow enabled their quantification in an automated and high-throughput manner, thereby offering new analytical value rather than claiming behavioral novelty.

The special resolution of RFID floor plate hardware is limited by the 5 x 5 cm grid size. The resolution is sufficient for proximity-based analyses, such as huddling or sleeping, where each grid typically contains only a few mice. However, higher resolution in RFID floor plate may be needed to capture fine-scale interactions like mating or grooming. It is important to emphasize that IntelliProfiler is purely a research workflow. Hardware specifications such as animal capacity or grid resolution are determined by the RFID board it is paired with and are not inherent limitations of the IntelliProfiler workflow itself. Notably, one RFID tag detached from a male mouse in the 16-mouse group, resulting in no movement data from Day 2 onward. As the untagged mouse remained in the cage, this may have slightly biased the social interaction data.

Despite its advantages, the RFID floor plate system (RFID technique) has a few limitations. First, the spatial resolution of RFID floor plate hardware is lower than video-based tracking systems like DLC^43,44^, as the current grid size is 5 cm x 5 cm, allowing two mice located in parallel within the same grid. Determining body orientation is also challenging when the animal remains within the same grid. Second, the RFID floor plate system has difficulty distinguishing detailed social contacts, such as sniffing, mounting, or huddling. Lastly, the current RFID system requires surgery for transponder implantation, and it is not yet possible to eliminate the invasive effects due to the size of commercially available transponders. These challenges should be addressed in future developments.

It has been reported that invasive interventions, such as intracranial surgery, can reduce continuous locomotor activity^45^. Although the RFID-tag implantation is considerably less invasive than neurosurgical procedures, we allowed a one-week recovery prior to behavioral assessments, following a previous study that implemented subcutaneous RFID transponder implantation in the abdominal region^46^. Another study reported that mice in pain may be socially avoided by conspecifics under naturalistic conditions, suggesting that healthy animals might actively avoid individuals experiencing disconfort^47^. In our experimental design, such avoidance responses were likely minimized, as all mice underwent RFID tag implantation, thereby standardized the physical condition and procedural experience across the group.

Importantly, IntelliProfiler is not a standalone system; it requires integration with the specific high-resolution RFID floor plate developed by Phenovance and currently incompatible with other hardware configurations. The RFID floor plate offers significant advantages in scalability and adaptability for high-throughput behavioral experiments in semi-naturalistic home-cage environments. This capability is especially valuable for neuroscience research, where detecting subtle behavioral differences is often critical. By enabling long-term, large-scale data collection with minimal human intervention, the RFID floor plate reduces experimenter bias and improves reproducibility.

The CCR metric, derived from IntelliProfiler workflow, provides a standardized, scalable measure of social interaction. When combined with additional behavioral parameters extracted via the workflow, the RFID floor plate enables a versatile approach to behavioral phenotyping. This integrated framework has broad applicability, ranging from basic neuroscience to translational preclinical research, with strong potential to advance studies on social behavior, neurodevelopmental disorders, and future pharmacological interventions.

Beyond mice, the RFID floor plate can be applied to other socially or nurturing species, such as rats, degus, and guinea pigs^48–50^. Furthermore, the combination of the RFID floor plate and IntelliProfiler could be used for quantitative assessment in models of neurodegenerative diseases, such as Alzheimer’s, by capturing spontaneous activity patterns. This approach offers a potential alternative to conventional behavioral testing protocols^42,51^.

## Conclusion

The combination of the RFID floor plate and IntelliProfiler workflow represents a significant advancement in animal behavior research, enabling continuous, detailed, and unbiased analysis of individual movements and social interactions within dynamic and complex group environments. In this study, we demonstrated its utility in uncovering the critical influence of aging, and an ASD model on behavioral patterns. As a robust and versatile platform, the RFID floor plate supports the study of behavior in more naturalistic home-cage settings, while the IntelliProfiler workflow provides opens new avenues for large-scale behavioral research and provides deeper insights into social dynamics.

## Methods

### Animals

C57BL/6J wild type (WT) mice at 7- or 52-week old were purchased from CLEA Japan, Inc., and maintained in the animal facility at Tohoku University Graduate School of Medicine. To generate aged father-derived offspring (AFO), 54–60-week-old male mice were mated with 8–12-week-old female mice at Tohoku University. For the AFO control group, 13–32-week-old male were mated with 10-12-week-old female mice at CLEA Japan, and their offspring were used as AFO controls. AFO mice were weaned at 4 weeks old and housed in littermate groups (2-5 animals per cage) until the start of experiments.

Following subcutaneous RFID tag implantation (see below), 7-week-old WT males were housed in groups of four per cage, 52-week-old aged WT males in pairs, and 7-week-old AFO males in groups of 2–5 per cage. All animals underwent a one-week recovery period prior to behavioral testing. Mice were housed in plastic cages (CL-0104-2; Clean S-TPX; 225 × 338 × 140 mm; CLEA Japan, Inc.) containing Aspen chip bedding (CLEA Japan) and water bottle sets (CK-200 K-11; CLEA Japan, CL-0904).

Unlike a previous study^52^ that employed long-term co-housing from the juvenile stage to minimize aggression, our design intentionally did not include an extended habituation period. Instead, adaptability to the social environment was evaluated following recovery from RFID implantation, with behavioral recording initiated at 8 or 53 weeks old. Although a short-term co-housing period was used for recovery, this was limited to littermate groups and was not intended as habituation to the full social group used in the behavioral assay.

Mice were maintained under standard laboratory conditions with a 12-hour light/dark cycle (lights on at 08:00 and lights off at 20:00), ambient temperature of 20–25 °C, and relative humidity of 40–60%. Animals were provided with the same standard chow (Lab MR Stock; Nihon Nosan Kogyo, Japan) and water *ad libitum*. Aspen chip bedding (CLEA Japan) was similarly used in all cages. All experimental procedures were approved by the Ethics Committee for Animal Experiments at the Tohoku University Graduate School of Medicine (approval number: 2021MdA-020-13).

### RFID implementation

Mice were deeply anesthetized using 5.0% isoflurane in an induction chamber (Shinano Manufacturing, Japan) and maintained under anesthesia with 2.0% isoflurane delivered via a facemask (Bio Research Center, Japan). Once fully anesthetized, each mouse was placed in a supine position, and the abdominal skin was gently lifted by pinching to facilitate subcutaneous access. An RFID tag (7 mm x 1.25 mm, Phenovance, Japan), preloaded in an injector, was inserted at a shallow angle from the caudal to cranial direction to ensure subcutaneous placement while avoiding penetration of the abdominal cavity. After fully inserting the needle tip beneath the skin, the RFID tag was slowly ejected, and gentle pressure was applied to stabilize the tag during needle withdrawal. Correct placement was immediately verified using a dual-band (HF/LF) RFID reader (Phenovance, Japan).

The implantation procedure employed in this study was minimally invasive and did not require an abdominal incision. Each implantation took approximately 2–3 minutes per animal, and mice typically recovered from anesthesia and resumed voluntary movement within a few minutes after the procedure. No obvious signs of distress or abnormal behavior were observed during the brief (∼5 minutes) post-surgical observation period. Mice were monitored during recover and allowed a one-week rest period prior to the commencement of behavioral recordings.

### Hardware

The antenaboard has been developed by Dr. Toshihiro Endo and his colleagues and is commercially available (eeeHive 2D, Phenovance LCC, Japan)^21^. The antenaboard consists of 96 antenna tiles (5 cm by 5 cm each) arranged in 12x8 grids (Fig. 1a). The home cage was 2000P cage (58 cm x 38 cm x 21.5 cm = width × depth × height, Tecniplast, Italy), equipped with a cage lid with food pellets, and three water bottles for long-term housing (Fig. 1a, b). The antenaboard and home cage were placed inside a soundproof box (Phenovance LCC, Japan). The light-dark cycle in the soundproof box and testing room was 12:12 hours (08:00-20:00, light; 20:00-08:00, dark). Mice were introduced at 8:00 in the light period on day 1.

### Data collection using RFID floor plate hardware

Data, including time, antenna ID, and transponder ID, were exported from the antenaboard hardware via Universal Serial Bus (USB) using TeraTerm software^53^ (Supplementary Fig. 1a, 1c).

### Software

R language (Version 4.2.2) was mainly used for preprocessing, data analysis, and visualization^54^. Python (Version 3.9.7), Prism 9, and Microsoft Excel (Version 16.86) were also used for preprocessing, data analysis, and visualization. R and Python packages required for the scripts are listed in Supplementary Table 1. All custom scripts are publicly available in a GitHub repository (see “Code Availability”).

### Preprocessing workflow

Raw data obtained from RFID floor plate hardware were processed into a format per second with time, antenna ID, transponder ID, and X-Y coordinates using the IntelliProfiler workflow (Supplementary Fig. 1b, 1d). In instances where multiple detections occurred within the same second, only the first X-Y coordinates were retained for analysis. Missing X-Y coordinates were interpolated by the immediately preceding value. These operations were performed using a custom R script, ‘IP_general.R’ (Supplementary Fig. 2).

### Importing RFID floor plate–derived data into the IntelliProfiler workflow

Positional information from the RFID floor plate was first logged as a plain text (.txt) file using the terminal software TeraTerm^53^. This raw log file served as the input to the workflow, initiated by a custom R script (IP_general.R) that parsed and reformatted the data for subsequent processing. This setup provided a structured transition from data acquisition to analysis, with downstream behavioral quantification performed through the IntelliProfiler workflow. Timestamps in the raw data were recorded using the computer clock *via* TeraTerm, which was synchronized with Japan Standard Time to ensure temporal consistency across all recorded sessions.

### Flow chart of RFID floor plate and IntelliProfiler workflow

The workflow is illustrated in Fig. 1c and Supplementary Fig. 2. Each stage: preprocessing and feature extraction, close contact ratio (CCR) computation, and advanced analysis is shown in the following **Methods** sections.

### Travel distance for each mouse

The Euclidean distance (D) of mouse movement was calculated from consecutive RFID positions at time *t* and *t+1* using the following formula:

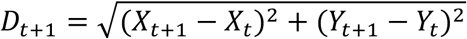

For each individual, the per-second distances were summed over time to yield a quantitative measure of physical activity. The total distance travelled over a given observation period *T* was calculated as:

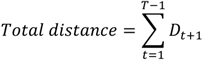

In this study, travel distance was evaluated across various time windows, such as 2 hours (*T* = 7,200 seconds) and 12 hours (*T* = 43,200 seconds), depending on the behavioral context or phase of interest (e.g., light vs. dark cycle).

This measure was pivotal for understanding the spatial dynamics of the subjects. Graphical representations of tracked movements, including 2D positional plots and time series plots of X and Y coordinates, were generated to visualize trajectories and movement patterns over time.

### Relative distance between two mice

The distances between different pairs of mice were calculated from their positions on the IntelliProfiler X–Y matrix, based on RFID floor plate data. For each pair of mice *i* and *j*, the inter-individual Euclidean distance at each time point *t* was computed as:

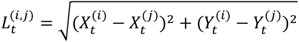

In addition, the total travel distance for each mouse was also calculated and plotted over the experimental period, providing a quantitative measure of activity levels. These calculations were performed using the custom R script “IP_general.R”.

### Spatial interaction analysis

Spatial relationships between paired mice on the X-Y plane were classified into four categories based on the calculated distance *L_t_*^(*i,j*)^ between the two mice (Fig. 3a). “**Same**” indicates that both mice occupied the same grid (*L* = 0 (cm)), suggesting immediate proximity. “**Close**” represents occupancy of adjacent grids (*L* = 5 or 5√2 (cm)), indicating close but distinct positions. “**Intermediate**” (*L* = 10, 5√5 or 10√2 (cm)) places one mouse on the outer perimeter of “**Close**,” range. “**Away**” (*L* > 10√2 (cm)) signifies that one mouse was outside of the “Intermediate” range, indicating substantial separation.

### Close contact ratio: defining social proximity

To quantify social interactions within a group of mice, we defined the Close Contact Ratio (CCR) as the cumulative percentage of time a pair of mice spent in the same grid (“**Same**”) or adjacent grids (“**Close**”).

CCR was computed over specified time windows (e.g., 2 hours or 12 hours) using the following formula:

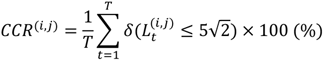

Here, *T* is the number of time points in the analysis window (e.g., *T* = 7,200 for 2 hours and *T* = 43,200 for 12 hours), and *δ*(·) is an indicator function returning 1 if the condition is satisfied or 0 otherwise. The threshold of 5√2 was chosen to reflect the maximal spatial extent for immediate or adjacent contact on the IntelliProfiler grid.

This metric provides a quantitative measure of sociality between two mice, indicating their tendency to remain in close physical proximity. Using Microsoft Excel and Prism 9 (GraphPad Software), we calculated CCR values as well as the proportionate distribution of all four distance categories (“**Same**,” “**Close**,” “**Intermediate**,” and “**Away**”) every 2 hours and separately for light and dark periods. All metrics were computed from output files generated by the “IP_general.R” script.

### Network analysis

Network analyses were performed for groups of four, eight, and 15/16 mice using Cytoscape 3.9.1^55^. For network construction, we prepared a tab-delimited summary table with three columns: Source, Target, and Edge betweenness. This table lists all mouse pairs along with their associated interaction strengths, calculated as:

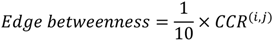

This downscaled CCR value was used to modulate edge thickness in Cytoscape visualizations^55^. To represent interaction topology, we applied the Edge-weighted Spring Embedded Layout, which facilitates intuitive interpretation of social interactions among mice based on their proximity metrics.

### Principal component analysis

Principal component analysis (PCA) was performed using Python 3 (Version 3.9.7) on four datasets derived from long-term behavioral recordings of mice in large groups during light and dark periods, focusing on relative activity levels and CCR. Three types of PCA visualizations were generated:

1. **PCA contribution vectors** displayed directly on the PCA plot (implemented in ‘IP_PCA.py’)
2. **K-means clustering analysis**, with the optimal number of clusters determined using the elbow method and silhouette method, applied to the PCA plot (implemented in ‘IP_Elbow_Silhouette.py’)
3. **PCA plots color-coded by gender and genotype** to facilitate group comparisons (implemented in ‘IP_PCA.py’).

All PCA computations and visualizations were performed using custom Python scripts (‘IP_PCA.py’ and ‘IP_Elbow_Silhouette.py’), which are available in the public GitHub repository (see **Code Availability**).

### Figure visualization

Graphical representations were primarily created using Prism 9 (GraphPad Software). Time-series line graphs and heat maps of travel distances or cumulative CCRs, as well as transitions in the proportionate of social distances (“**Same**,” “**Close**,” “**Intermediate**,” and “**Away**”) were generated. Pie charts illustrating the distribution of social distances were also prepared. PCA plots were created in Python 3, and the social network graphs for the groups were constructed using Cytoscape version 3.9.1^55^.

### Statistical analysis

Statistical analyses were conducted with Prism version 9 (GraphPad). Two-way ANOVA followed by Holm-Sidak’s multiple comparisons test was used to assess differences between groups; part these analyses has been presented in part in a previous article^56^. Data in bar or line graphs are presented as mean ± SD, with statistical significance indicated as *p*\* < 0.05, *p*\*\*<0.01, *p*\*\*\*<0.001. Exact *p*-values, *F* values, and degrees of freedom for all results are provided in Supplementary Table 2. All experiments were performed once (single repetition).

## Data Availability

The behavioral data generated are provided in the **Supplementary Information**.

## Code Availability

All code and scripts used for data analysis are available on GitHub.

- Repository URL: https://github.com/ShoheiOchi/IntelliProfiler

## Acknowledgements

The authors are grateful to Dr. Ken-ichiro Tsutsui for temporally providing the behavioral analysis environment, and Kentaro Abe for advising on the setup of the behavioral analysis room. We also thank Dr. Toshihiro Endo and his colleagues for developing a RFID floor plate used in this study, especially, Dr. Endo for technical advices. This research was funded by the JSPS KAKENHI under grant number 24H01419 (to S.O.) and 24K02203 (to N.O.), AMED under grant number JP21wm0425003 (to N.O.) JP24wm0225044 (to S.O.). Takeda Science Foundation (to S.O.) and FY2024 Tohoku University Graduate School of Medicine Grant-in-Aid for Joint Research by Young Researchers (to S.O.).

## Contributions

Conceptualization, S.O., H.I. and N.O.; behavior analysis, S.O.; data process, S.O. and H.I.; code preparation, S.O. and H.I.; figure preparation, S.O.; writing—review and editing, S.O., H.I. and N.O.; supervision, N.O.; project administration, S.O. and N.O.; funding acquisition, S.O., and N.O. All authors have read and agreed to the published version of the manuscript.

