## Supplementary information for "IntelliProfiler: a research workflow for analyzing multiple animals with a high-resolution home-cage RFID system"

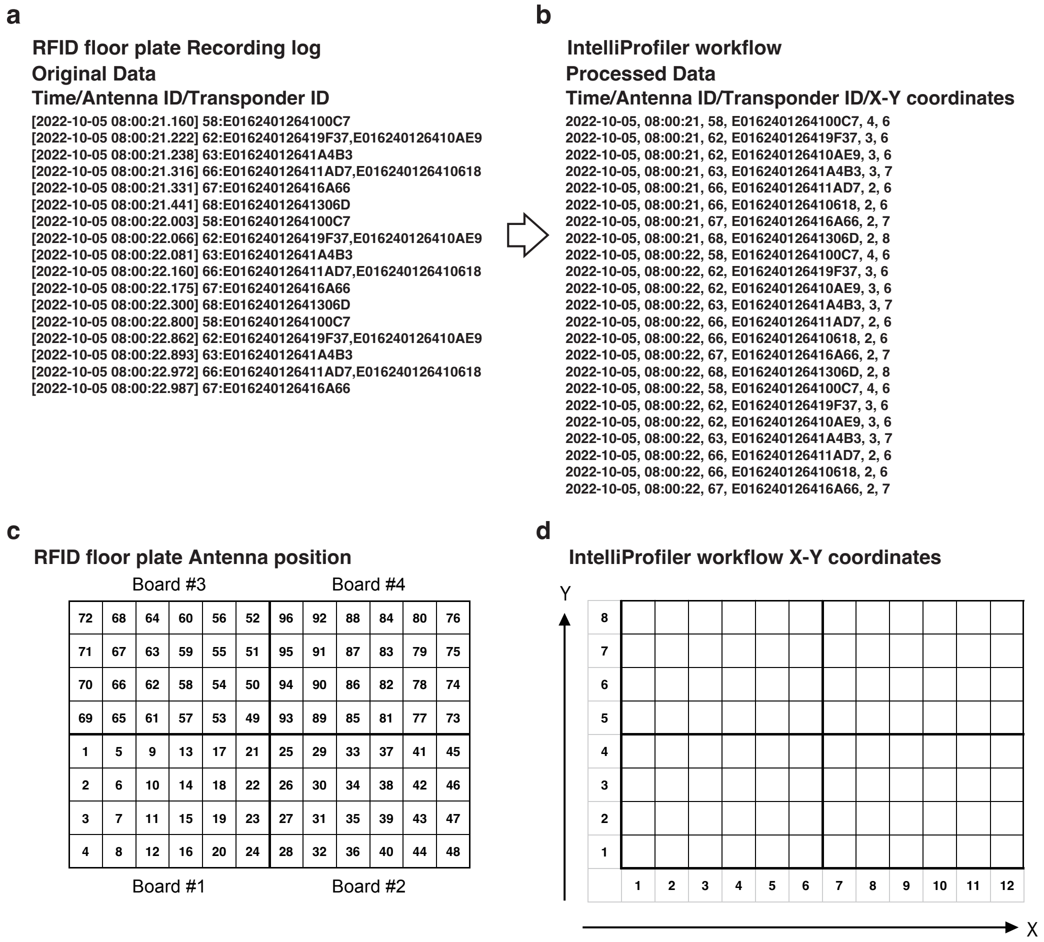


**Supplementary Fig. 1: RFID floor-plate raw data and data processing *via* the IntelliProfiler workflow.**

**a** Original data capturing Time, Antenna ID, and Transponder ID from the RFID floor plate. **b** Processed data generated *via* the IntelliProfiler workflow, refined to include Time, Antenna ID, Transponder ID, and XY coordinates. Decimal values are removed from the time data. If multiple individuals are detected on the same antenna ID simultaneously, their information is separated into different rows. **c** Layout and antenna numbering of the four RFID floor plates. **d** Conversion of antenna positions to X-Y coordinates across the four RFID floor plates, mapped to a X-Y coordinate of X1–12 and Y1–8 *via* the IntelliProfiler workflow.

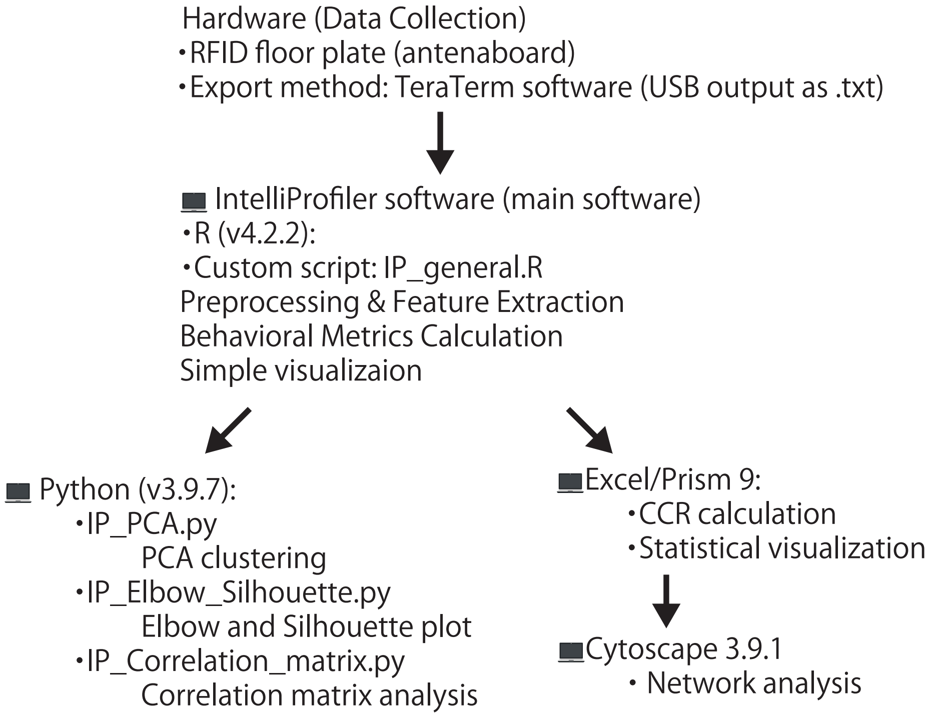


**Supplementary Fig. 2: Flow chart of IntelliProifler research workflow.**
The IntelliProfiler workflow consists of four main stages for analyzing group-housed mouse behavior using RFID-based position data. Data are collected from an RFID floor plate and preprocessed using a custom R script (IP_general.R) to interpolate missing values and assign X-Y coordinates. Behavioral metrics, including travel distance and inter-individual distance, are calculated in R, while close contact ratio (CCR) is computed in Microsoft Excel. Visualizations such as 2D plots, PCA, correlation matrices, and social networks are generated using R, Python scripts (IP_PCA.py, IP_Elbow_Silhouette.py, IP_Correlation_matrix.py), Excel, and Cytoscape. All scripts used in the IntelliProfiler workflow is available in the public GitHub repository (see "Code Availability").

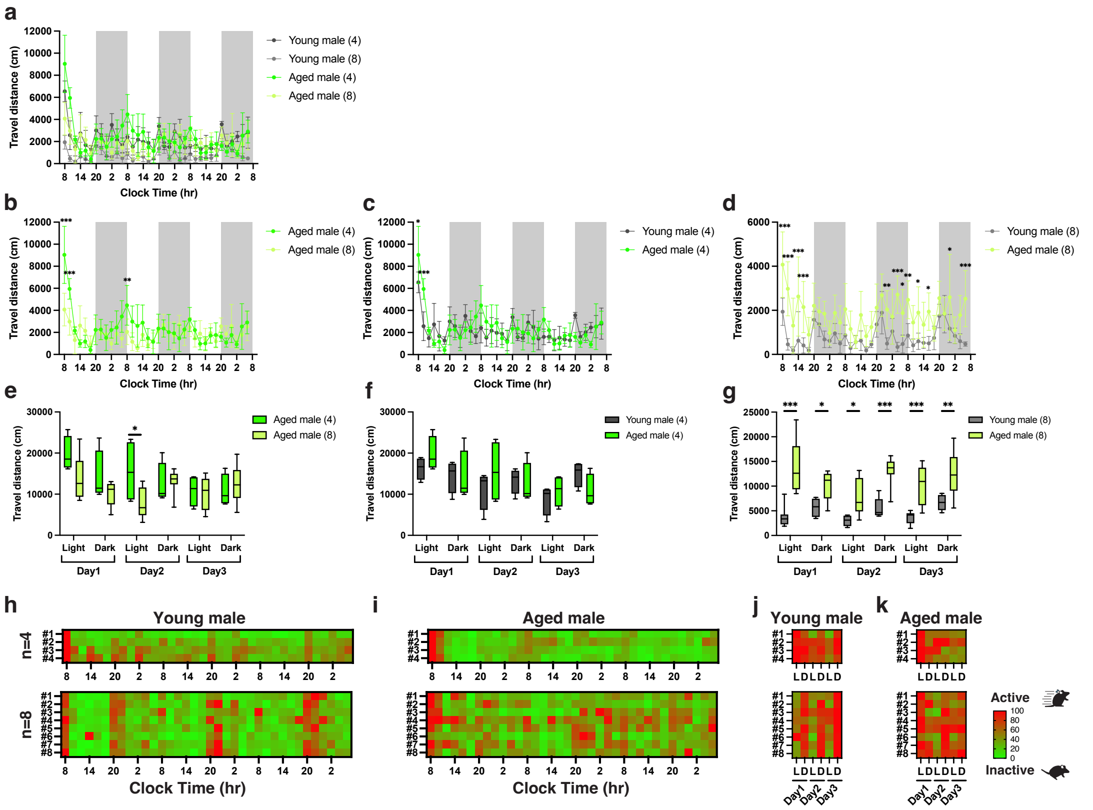


**Supplementary Fig. 3: Activity analysis of group-housed aged male mice using the IntelliProfiler workflow.**

**a-g** Cumulative travel distances measured in 2-hour intervals (**a-d**) and over 12-hour intervals (**e-g**). **b, e** Analysis of locomotor activity based on group size, comparing groups of four and eight aged mice. **c, d, f, g** Analysis of aging effects in locomotor activity, comparing groups of four young and aged male mice (**c, f**), groups of eight young and aged male mice (**d, g**). **h-k** Heatmaps showing relative locomotor activity over 2-hour intervals (**h, i**) and 12-hour intervals (**j, k**) for groups of four and eight mice. Values in the bar or line graphs are presented as mean ± SD, with box plots displaying min/max whiskers. Statistical analysis was conducted using Holm-Sidak method for multiple comparisons. Significance levels are indicated as *$p < 0.05$, **$p < 0.01$, and ***$p < 0.001$. $p = 0.338$ (**c**), $p < 0.001$ (**d**), $p = 0.585$ (**f**), $p < 0.001$ (**g**).

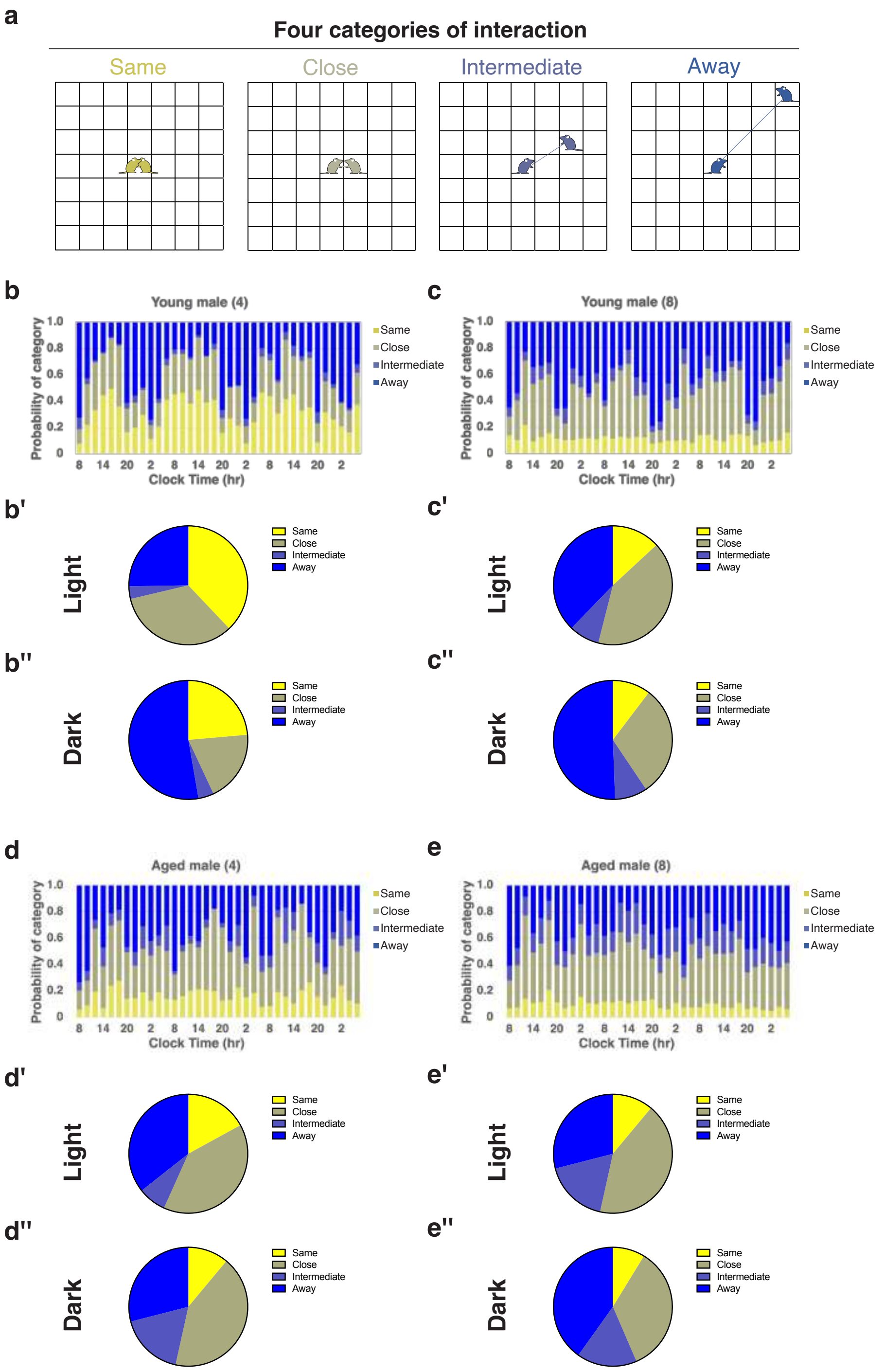


**Supplementary Fig. 4: The spatial relationships among group-housed aged male mice using the IntelliProfiler workflow.**

**a** Definition of the spatial relationships between two individual mice. When another mouse is in the same grid as the focal mouse (both depicted in yellow), the relationship is defined as “Same.” If the other mouse is in the surrounding grids (depicted in olive), it is defined as “Close.” If the other mouse is outside the surrounding grids (depicted in light indigo), it is defined as “Intermediate.” If the other mouse is beyond the “Intermediate” zone (depicted in blue), the relationship is categorized as “Away.” **b-e** The 2-hour bin transition of proportions based on “Same,” “Close,” “Intermediate,” and “Away” for groups of four young male mice (**b**), eight young male mice (**c**), four aged male mice (**d**), eight aged male mice (**e**). **b-e** The average proportion of distance categories during the light period and the dark period is shown for each group. Chi-square $p$-values comparing the proportions between light and dark periods, $p < 0.001$ (**b**, four young males), $p = 0.256$ (**c**, eight young males), $p = 0.836$ (**d**, four aged males), $p = 0.439$ (**e**, eight aged males). The $p$-values of chi-square for the group size, $p < 0.001$ (**b, c**, young males in the light phase), $p = 0.02$ (**b, c**, young males in the dark phase), $p = 0.115$ (**d, e**, aged males in the light phase), $p = 0.477$ (**d, e**, aged males in the dark phase).

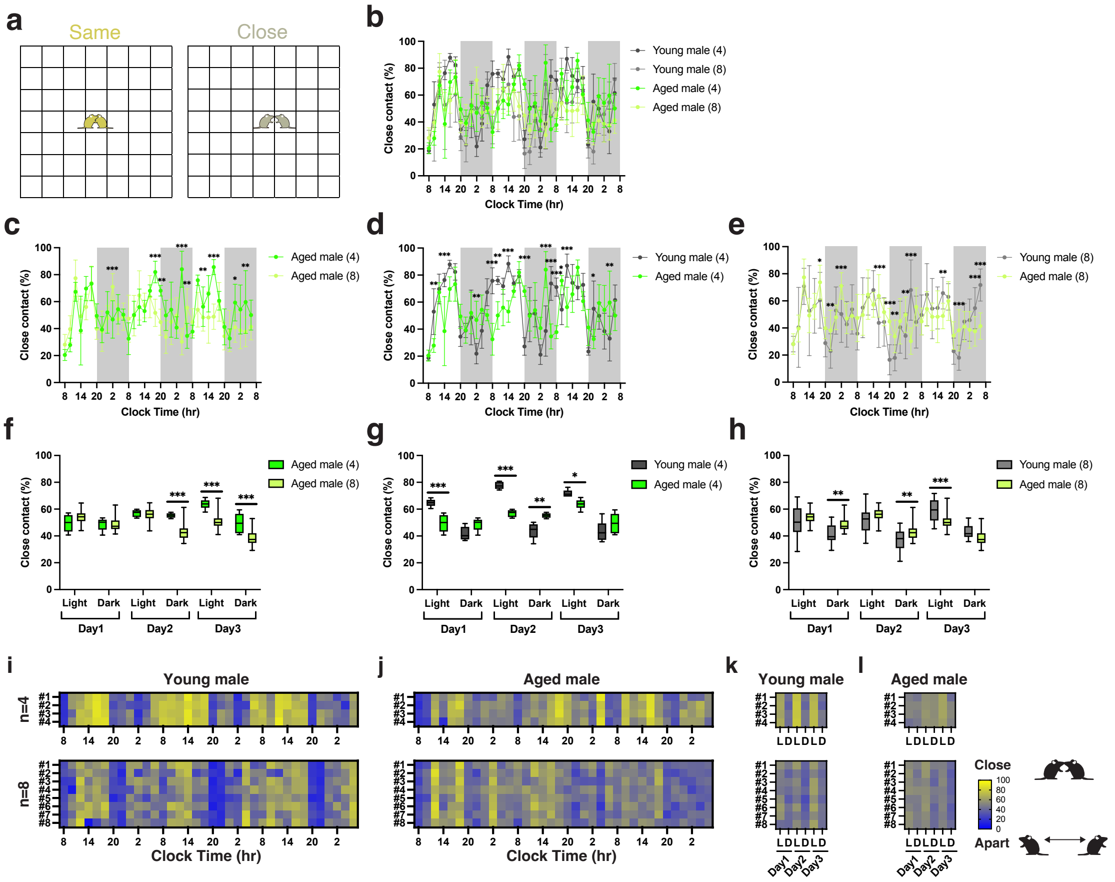


**Supplementary Fig. 5: Close contact between two individuals among group-housed aged male mice analyzed using the IntelliProfiler workflow.**

**a** Definition of close contact, where two mice either in either “Same” (depicted in yellow) or in adjacent grids, classified as “Close” (depicted in olive). **b-h** Average Close Contact Ratio (CCR) measured in 2-hour intervals (**b-e**) and 12-hour intervals (**f-h**). **c, f** Analysis of CCR based on group size, comparing groups of four and eight young and aged males. **d, e, g, h** Analysis of CCR based on aging effects, comparing groups of four young and aged male mice (**d, g**), groups of eight young and aged male mice (**e, h**). **i-l** Heatmaps showing the time course of CCR over 2-hour intervals (**i, j**) and 12-hour intervals (**k, l**) for groups of four and eight mice. Values in the bar or line graphs are presented as mean ± SD, with box plot showing min/max whiskers. Statistical analysis was conducted using Holm-Sidak method for multiple comparisons. Significance levels are indicated as *$p < 0.05$, **$p < 0.01$, and ***$p < 0.001$. $p < 0.001$ (**c**), $p = 0.338$ (**d**), $p = 0.072$ (**e**), $p < 0.001$ (**f**), $p = 0.007$ (**g**), $p = 0.109$ (**h**).

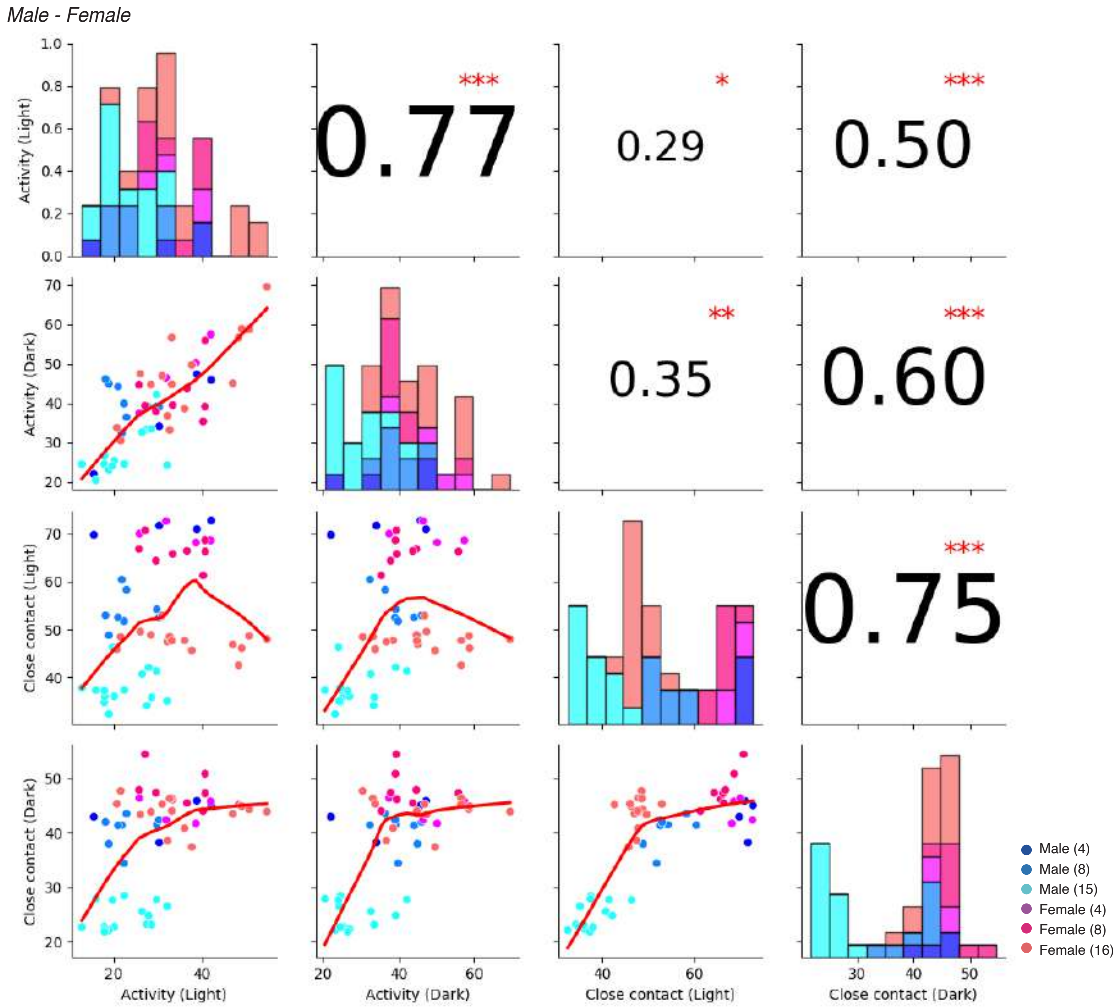


**Supplementary Fig. 6: Correlation matrix and histograms comparing relative activity and close contact between males and females.**

This figure presents a correlation matrix and histograms comparing relative locomotor activity and Close Contact Ratio (CCR) between male and female mice. The diagonal elements represent histograms of each behavior parameter: relative activity during the light period (Light), relative activity during the dark period (Dark), CCR during the light period (Light), and CCR during the dark period (Dark). The lower triangular plots show the scatter plots with fitted regression lines, while the upper triangular plots display the Pearson correlation coefficients between each pair of variables. *$p < 0.05$, **$p < 0.01$, ***$p < 0.001$. Circular points are color-coded to represent different groups: four, eight, 15-16 males and females.

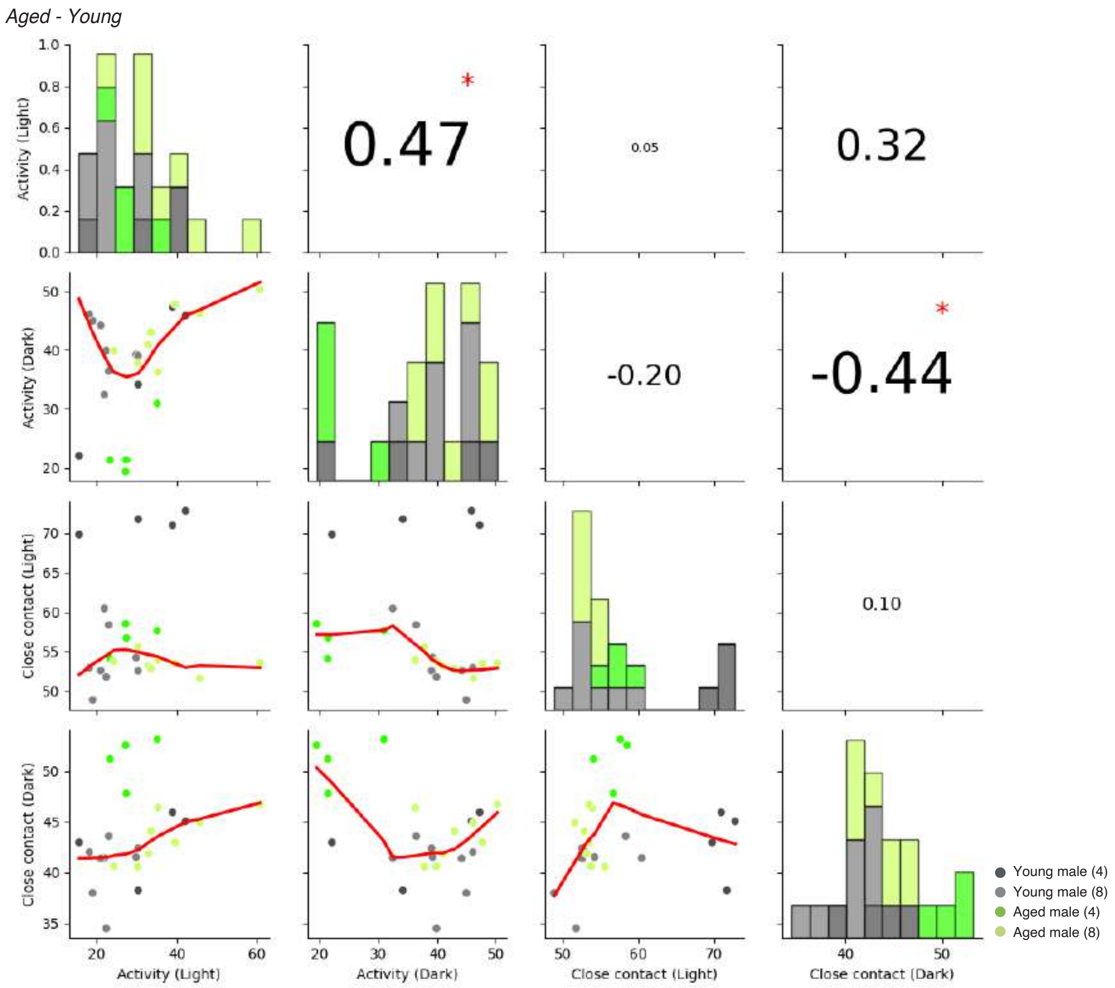


**Supplementary Fig. 7: Correlation matrix and histograms comparing relative activity and close contact between young and aged males.**

This figure presents a correlation matrix and histograms comparing relative locomotor activity and the Close Contact Ratio (CCR) between young and aged mice. The diagonal elements represent histograms of each behavior parameter: relative activity during the light period (Light), relative activity during the dark period (Dark), CCR during the light period (Light), and CCR during the dark period (Dark). The lower triangular plots show the scatter plots with fitted regression lines, while the upper triangular plots display the Pearson correlation coefficients between each pair of variables. *$p < 0.05$. Circular points are color-coded to represent different groups: four, and eight young and aged males.

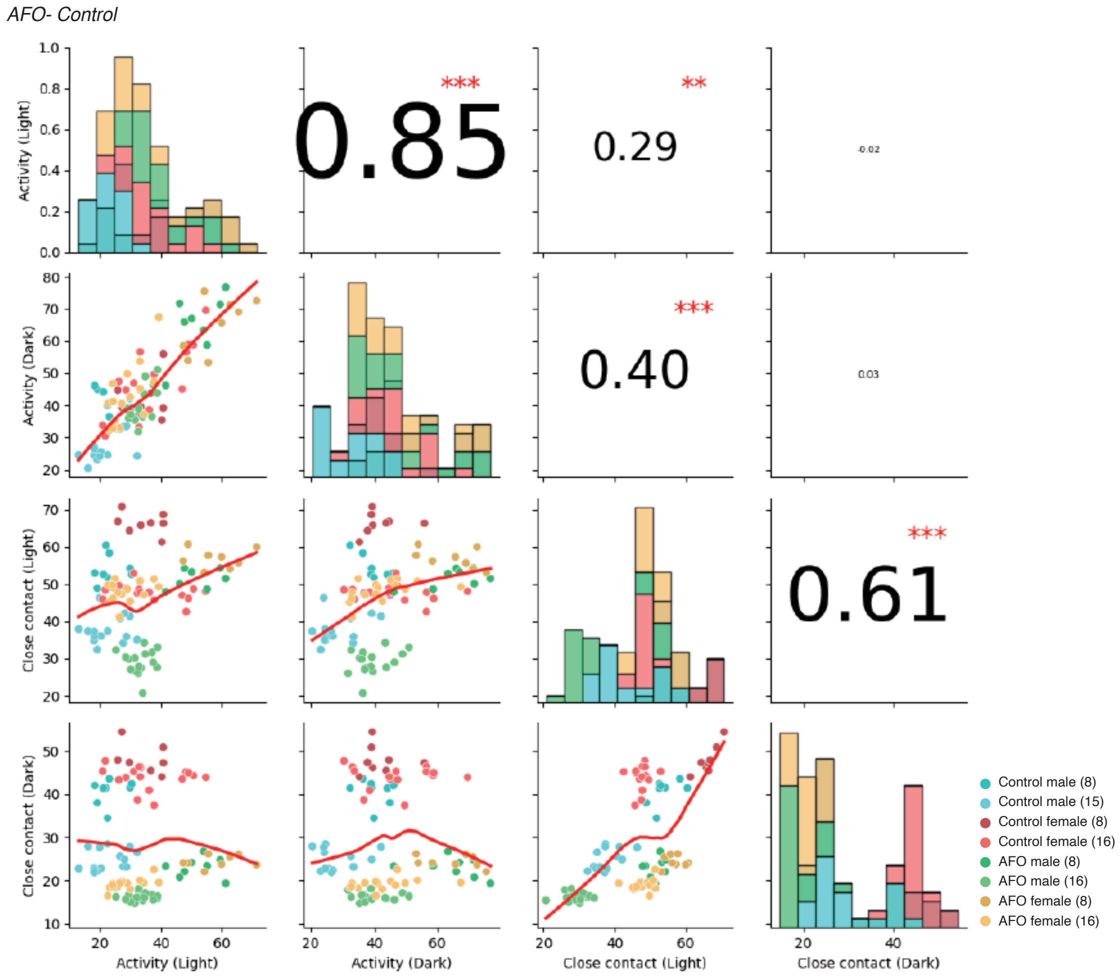


**Supplementary Fig. 8: Correlation matrix and histograms comparing relative activity and close contact between AFO and control males and females.**

This figure presents a correlation matrix and histograms comparing relative locomotor activity and the Close Contact Ratio (CCR) between AFO and control mice. The diagonal elements represent histograms of each behavior parameter: relative activity during the light period (Light), relative activity during the dark period (Dark), CCR during the light period (Light), and CCR during the dark period (Dark). The lower triangular plots show the scatter plots with fitted regression lines, while the upper triangular plots display the Pearson correlation coefficients between each pair of variables. $^{**}p < 0.01$, $^{***}p < 0.001$. Circular points are color-coded to represent different groups: eight and 16 AFO and control males and females.

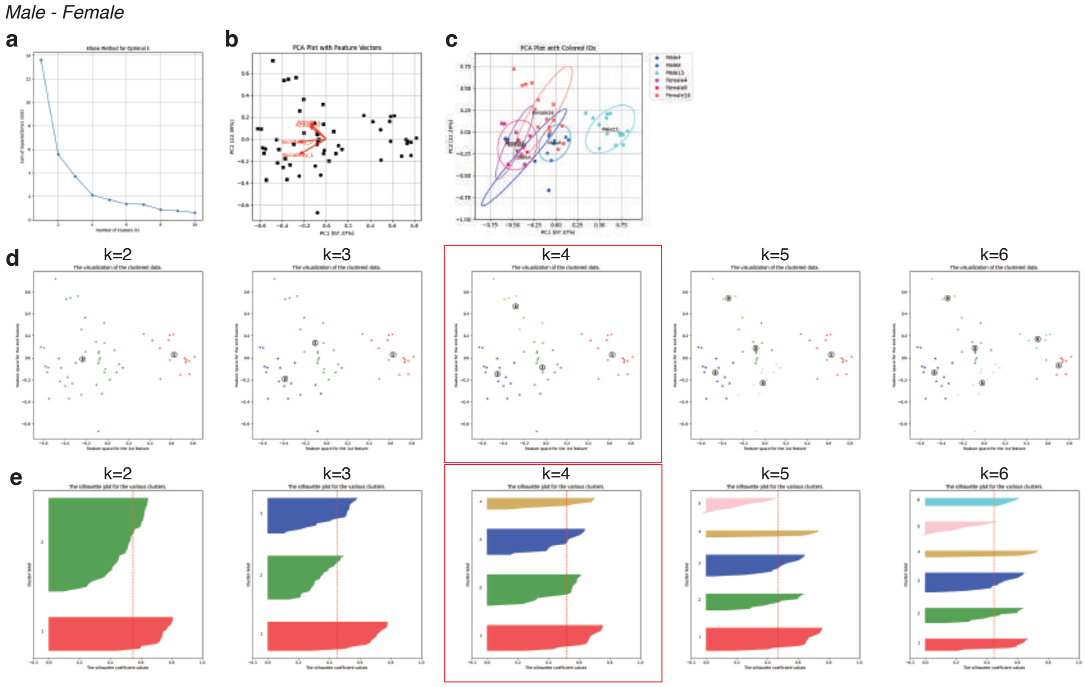


**Supplementary Fig. 9: Determination of the optimal cluster number for K-means clustering using the Elbow method and Silhouette analysis in males and females using the IntelliProfiler workflow.**

This figure illustrates the determination of the optimal cluster number for K-means using the Elbow method and Silhouette analysis in groups of four, eight, and 15/16 male and female mice (**a-e**). **a** The Elbow method and silhouette analysis are used to estimate the appropriate number of clusters (k). The plot displays the sum of squared errors (SSE) against the number of clusters (k). **b** Analysis of feature vectors for Close Contact Ratio (CCR) and relative activity during the light and dark periods is shown in PCA plots. **c** PCA plots displaying the distribution of individual points, which are color-coded. Confidence ellipses (95%) are overlaid to visualize the group-level variance in PCA space. **d** K-means clustering results with different cluster numbers (k=2 to k=6). Each panel illustrates the distribution of the individual points in PCA space along with their respective cluster assignments. **e** Silhouette analysis plots corresponding to the K-means clustering results shown in panels **d**. The silhouette coefficient values are plotted for each data point, with the red dashed line representing the average silhouette score for each k. The optimal cluster numbers are highlighted with red lines (**d, e**). A summarized PCA analysis is presented in **Fig. 8a-c**.

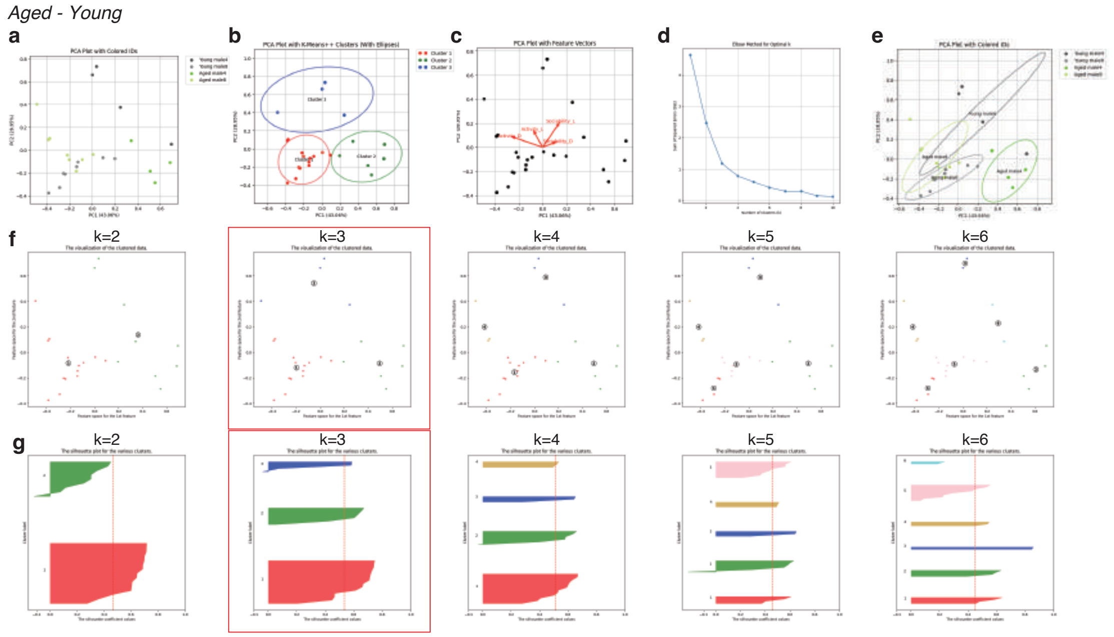


**Supplementary Fig. 10: Principal component analysis of group behavior in aged males using the IntelliProfiler workflow.**

Principal component analysis (PCA) of group behavior in four, eight aged and young male mice. **b** K-means clustering in PCA plots. **c** Analysis of feature vectors, focusing on Close Contact Ratio (CCR) and relative activity during the light and dark periods. **d** The Elbow method and silhouette analysis are used to estimate the appropriate number of clusters (k). The plot displays the sum of squared errors (SSE) against the number of clusters (k). **e** PCA plots displaying the distribution of individual points, which are color-coded. Confidence ellipses (95%) are overlaid to visualize the group-level variance in PCA space. **f** K-means clustering results with different cluster numbers (k=2 to k=6). Each panel illustrates the distribution of the individual points in PCA space along with their respective cluster assignments. **g** Silhouette analysis plots corresponding to the K-means clustering results shown in panels **f**. The silhouette coefficient values are plotted for each data point, with the red dashed line representing the average silhouette score for each k. The optimal cluster numbers are highlighted with red lines (**f, g**). A summarized PCA analysis is presented in **a-c**.

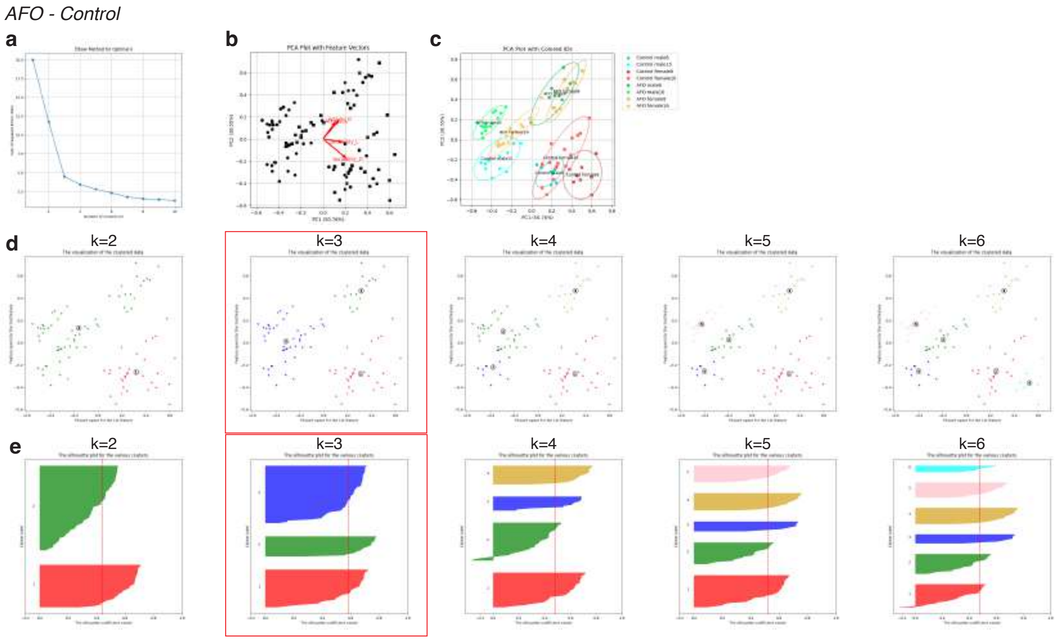


**Supplementary Fig. 11. Determination of the optimal cluster number for K-means clustering using the Elbow method and Silhouette analysis in AFO and control males and females using the IntelliProfiler workflow.**
This figure illustrates the determination of the optimal cluster number for K-means using the Elbow method and Silhouette analysis in groups of eight, and 15/16 control and AFO male and female mice (**a-e**). **a** The Elbow method and silhouette analysis are used to estimate the appropriate number of clusters (k). The plot displays the sum of squared errors (SSE) against the number of clusters (k). **b** Analysis of feature vectors for Close Contact Ratio (CCR) and relative activity during the light and dark periods is shown in PCA plots. **c** PCA plots displaying the distribution of individual points, which are color-coded. Confidence ellipses (95%) are overlaid to visualize the group-level variance in PCA space. **d** K-means clustering results with different cluster numbers (k=2 to k=6). Each panel illustrates the distribution of the individual points in PCA space along with their respective cluster assignments. **e** Silhouette analysis plots corresponding to the K-means clustering results shown in panels **d**. The silhouette coefficient values are plotted for each data point, with the red dashed line representing the average silhouette score for each k. The optimal cluster numbers are highlighted with red lines (**d, e**). A summarized PCA analysis is presented in **Fig. 8d-f**.

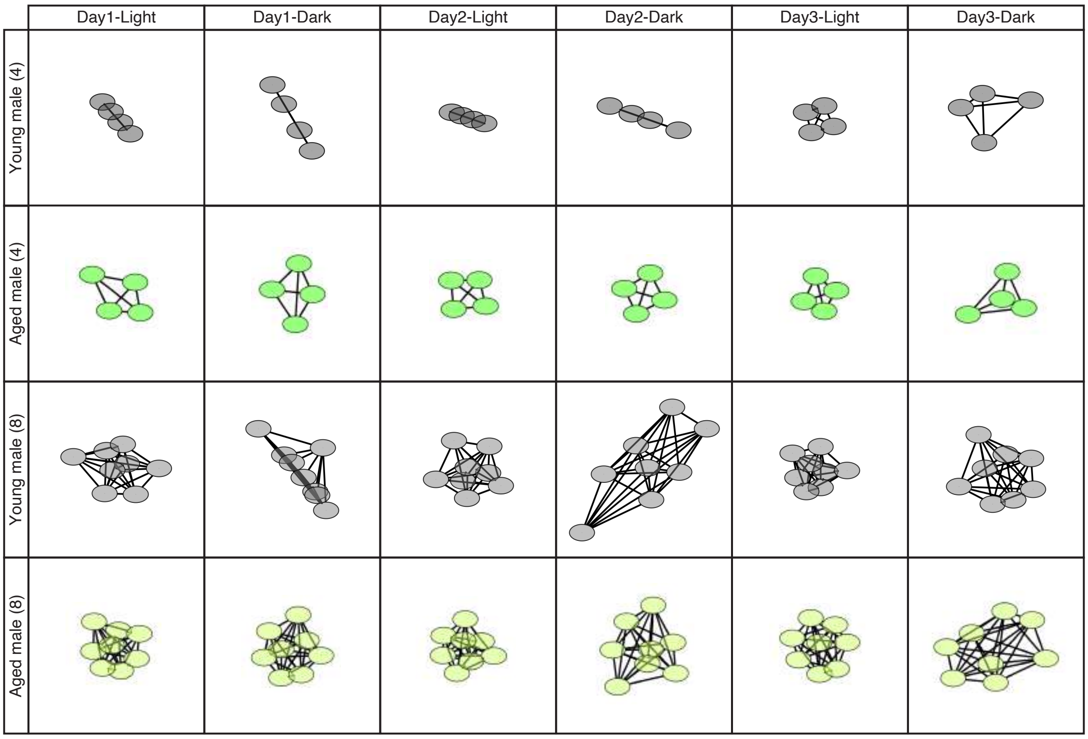


**Supplementary Fig. 12: Network analysis of group interaction in young and aged males using the IntelliProfiler workflow.**

Network analysis of social interactions during the light period on day 1 and the dark period on day 3 in groups of four and eight young and aged males. Each circle represents an individual mouse, and the proximity of the lines indicates the relative social distance between two individuals. The network graph was generated using Cytoscape.

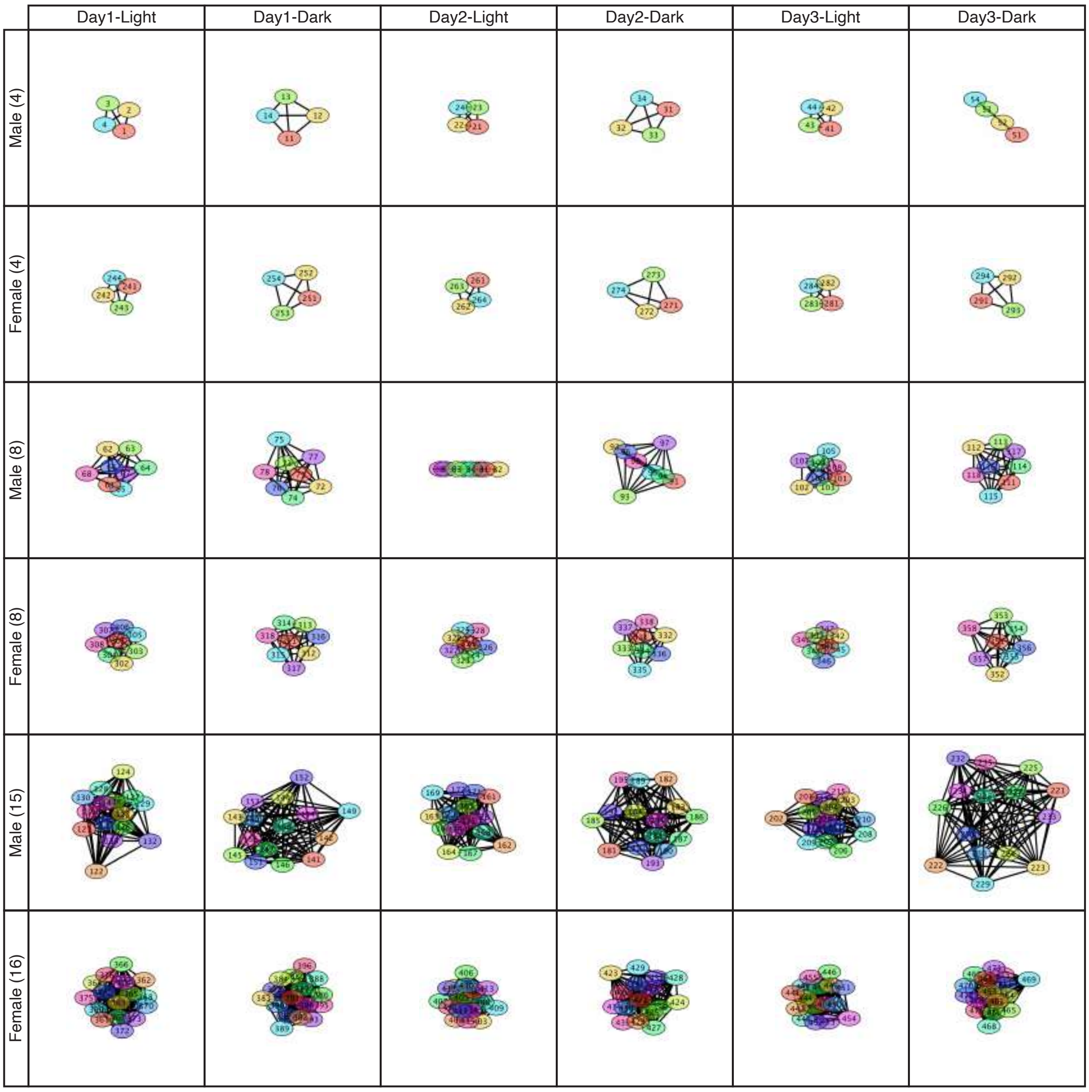


**Supplementary Fig. 13: Network analysis of social interaction in the color-coded male and female individuals.**

This figure presents a network analysis of social interactions during the light period on day 1 and the dark period on day 3 in groups of four, eight, and 15/16 male and female mice. Nodes represents individual mice, each assigned a unique color codes, while edges reflect the relative social contacts between two individuals. The network graph was created using Cytoscape. The network diagrams, shown in **Fig. 9**, utilize specific color codes for each group, highlighting the differences and similarities in social interactions among the various groups.

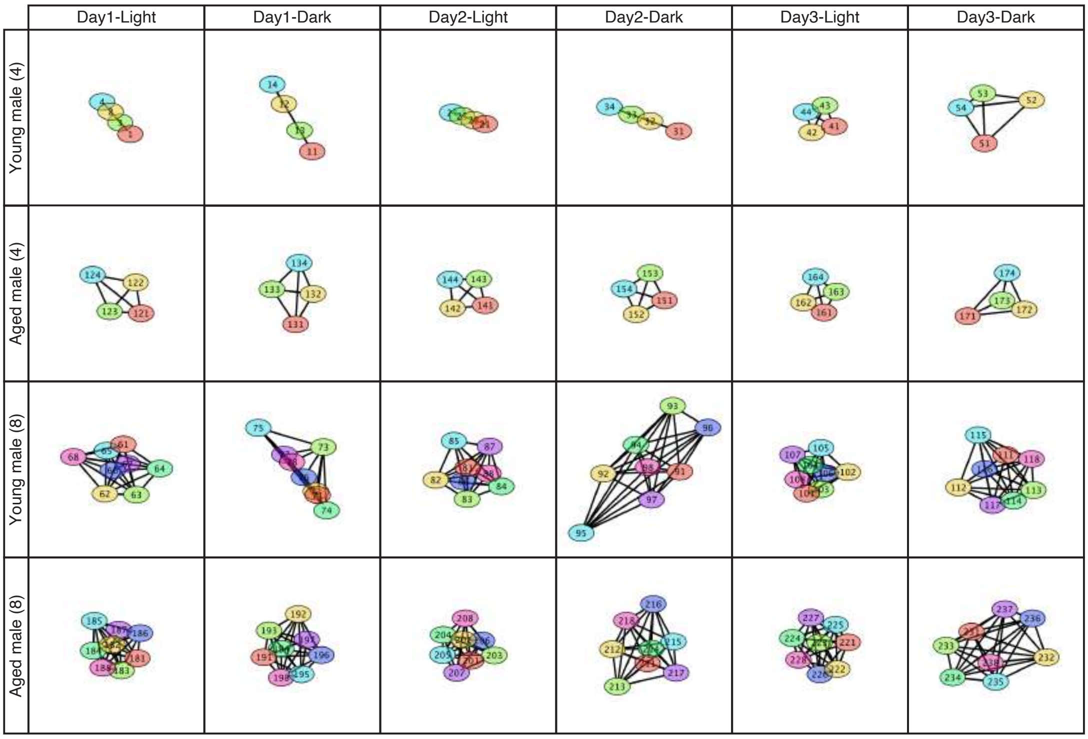


**Supplementary Fig. 14: Network analysis of social interaction in the color-coded young and aged male individuals.**

This figure presents a network analysis of social interactions during the light period on day 1 and the dark period on day 3 in groups of four and eight young and aged male mice. Nodes represents individual mice, each assigned a unique color code, while edges reflect the relative social contacts between two individuals. The network graph is created using Cytoscape. The network diagrams, displayed in **Supplementary Fig. 11**, utilize specific color codes for each group, highlighting the differences and similarities in social interactions among the various groups.

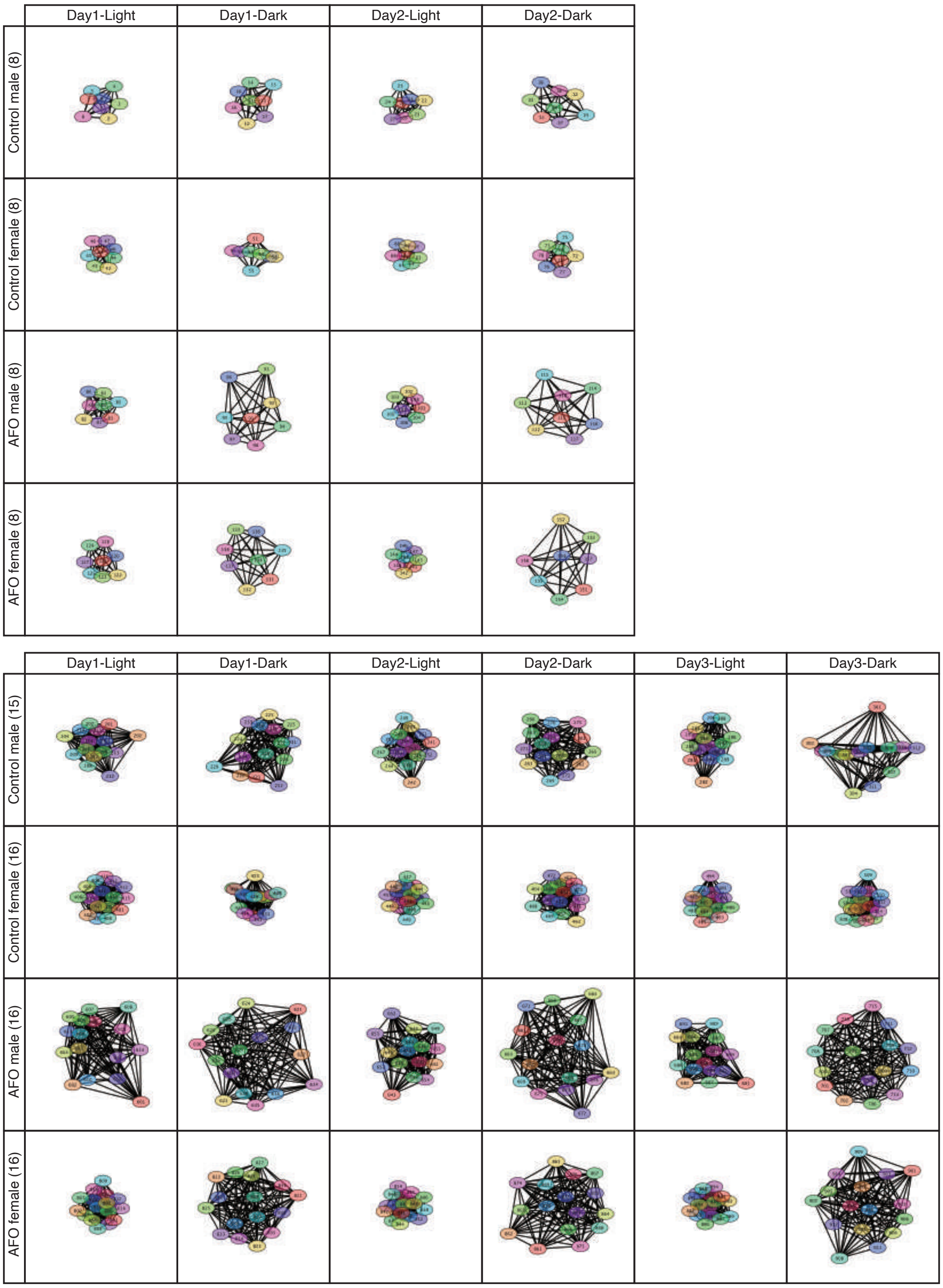


**Supplementary Fig. 15: Network analysis of social interaction in the color-coded male and female AFO and control individuals.**

This figure presents a network analysis of social interactions during the light period on day 1 and the dark period on day 2 in groups of eight AFO and control mice and during the light period on day 1 and the dark period on day 3 in groups of 16 AFO and control mice. Nodes represents individual mice, each assigned a unique color code, while edges reflect the relative social contacts between two individuals. The network graph is created using Cytoscape. The network diagrams, displayed in **Fig. 10**, utilize specific color codes for each group, highlighting the differences and similarities in social interactions among the various groups.

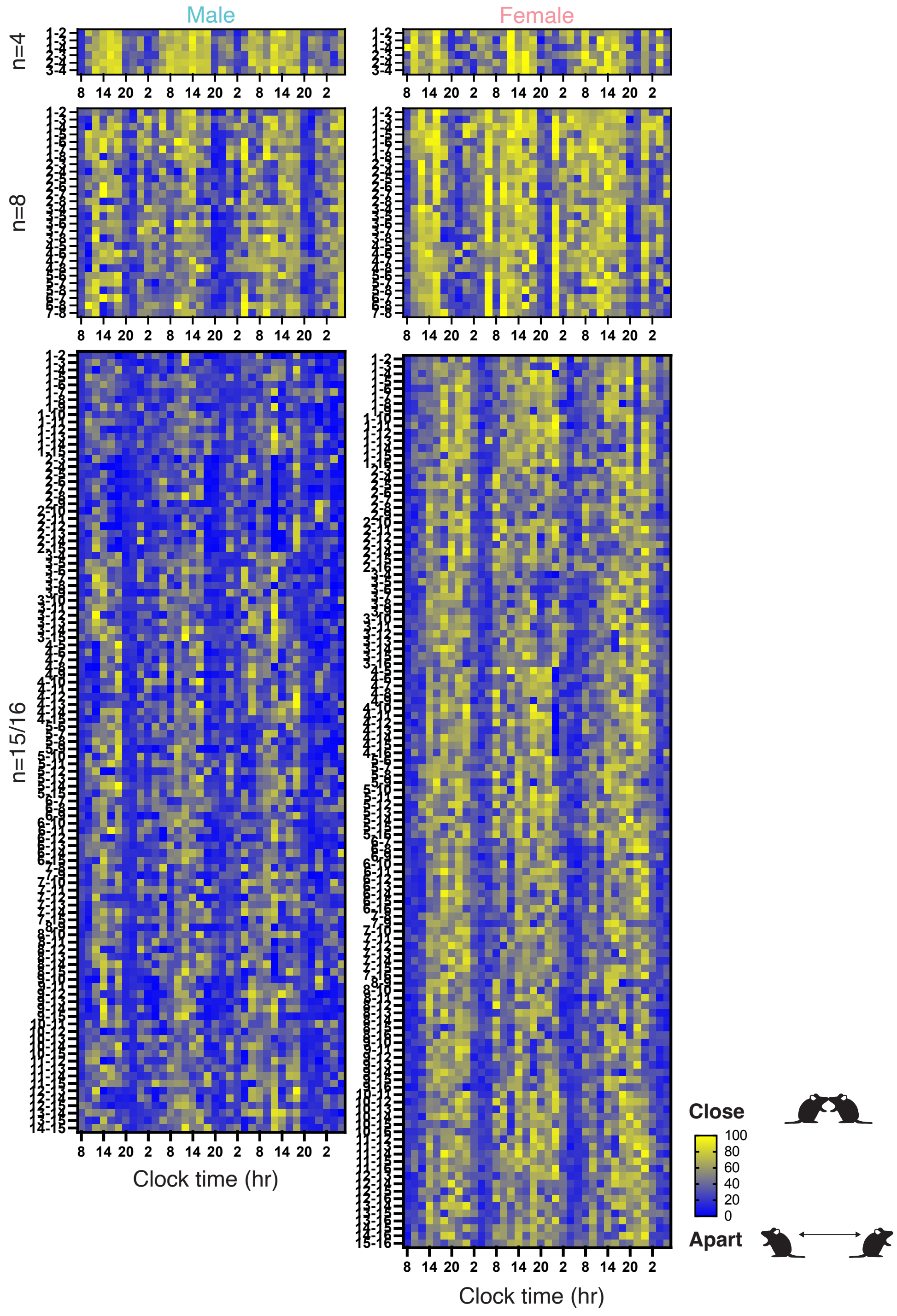


**Supplementary Fig. 16: Analysis of pairwise social interaction in males and females using the IntelliProfiller workflow.**

This heatmap illustrates the time course of Close Contact Ratio (CCR) between two individuals every 2 hours within groups of four, eight, and 15/16 male and female mice.

Hypothetical mouse ID #X and #Y are designated, with the two individuals represented as X-Y to facilitate analysis. The total number of pair combinations is calculated using the mathematical formula ${}_xC_2$, resulting in six combinations for groups of four males and females, 28 for groups of eight males and females, 105 for a group of 15 males, and 120 for a group of 16 females.

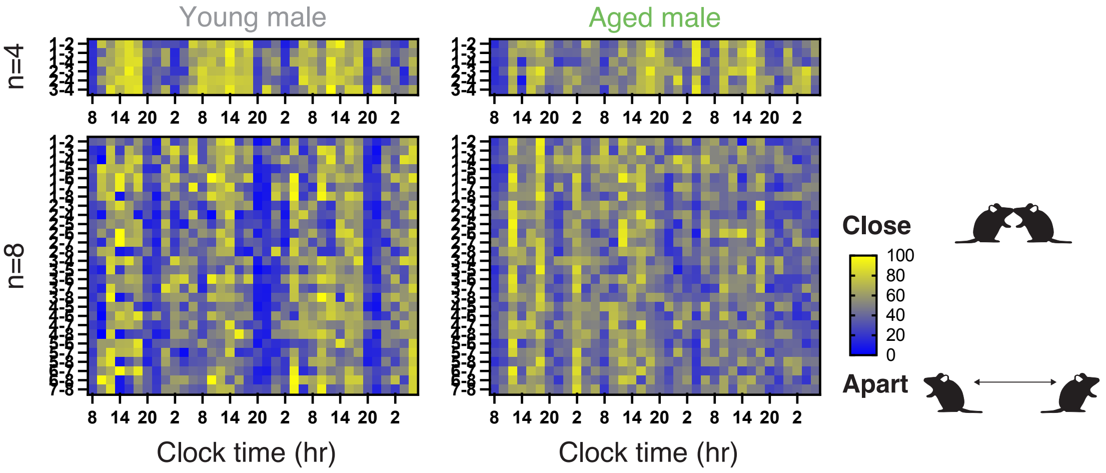


**Supplementary Fig. 17: Analysis of pairwise social interaction in young and aged males using the IntelliProfiller workflow.**

This heatmap illustrates the time course of Close Contact Ratio (CCR) between two individuals every 2 hours within groups of four and eight young and aged male mice. Hypothetical mouse ID #X and #Y are designated, with the two individuals represented as X-Y to facilitate analysis. The total number of pair combinations is calculated using the mathematical formula $_{x}C_{2}$, resulting in six combinations for groups of four young and aged males, 28 for groups of eight young and aged males.

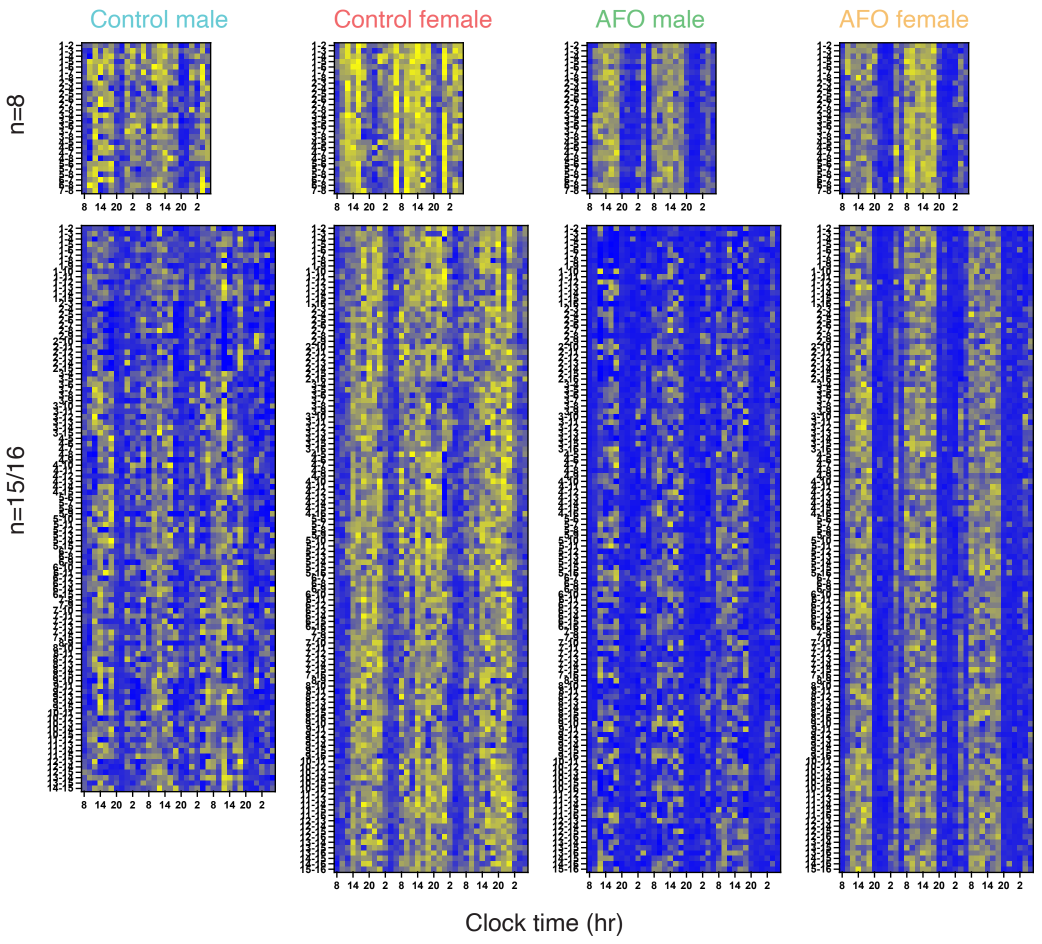


**Supplementary Fig. 18: Analysis of pairwise social interaction in AFO and control males and females using the IntelliProfiller workflow.**

This heatmap illustrates the time course of Close Contact Ratio (CCR) between two individuals every 2 hours within groups of eight and 15/16 AFO and control male and female mice. Hypothetical mouse ID #X and #Y are designated, with the two individuals represented as X-Y to facilitate analysis. The total number of pair combinations is calculated using the mathematical formula ${}_xC_2$, resulting in six combinations for groups of 28 for groups of eight AFO and control males and females, 105 for a group of 15 control males, and 120 for a group of 16 AFO males and control and AFO females.

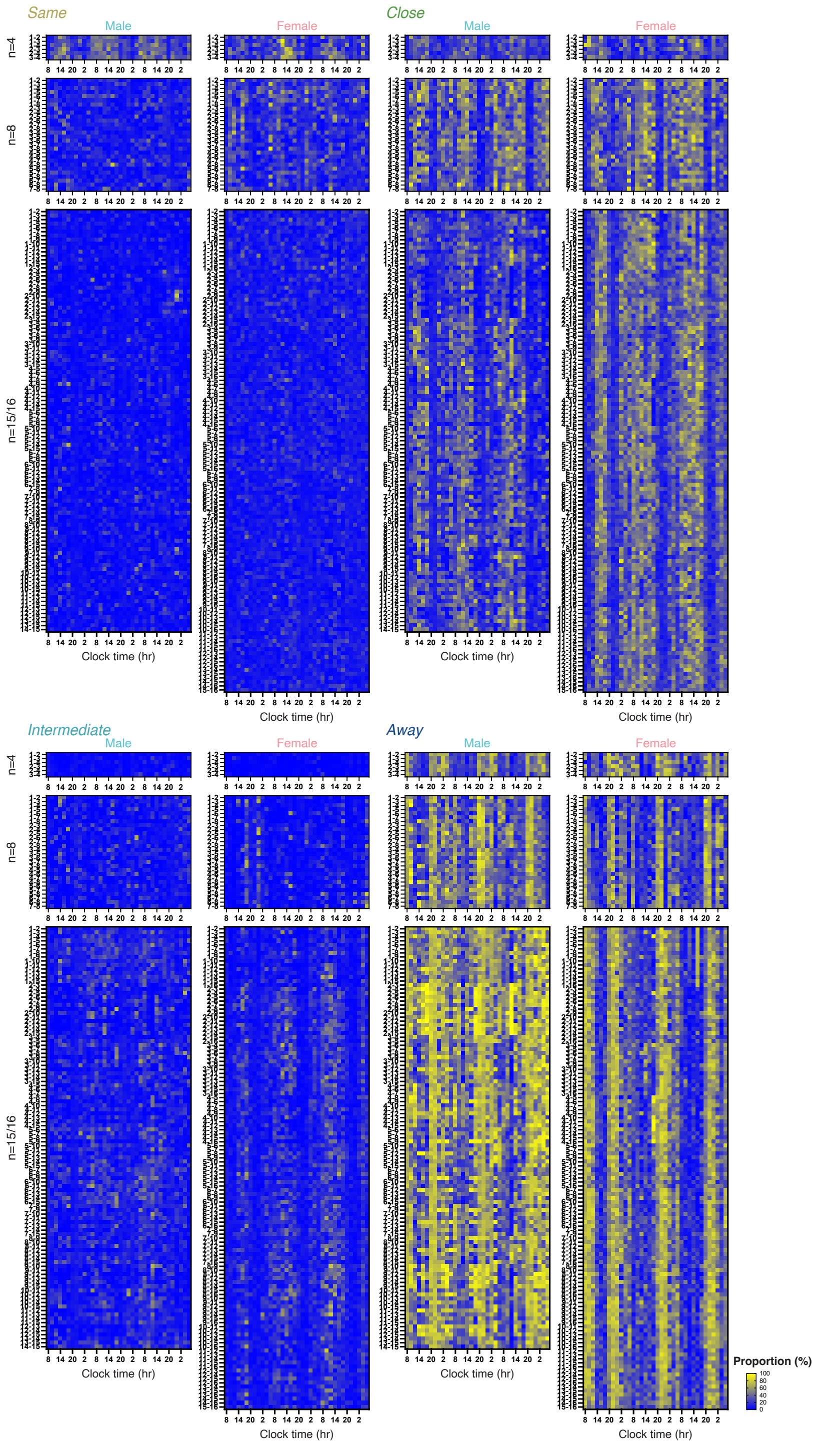


**Supplementary Fig. 19: Analysis of pairwise spatial relationships in male and female individuals using the IntelliProfiller workflow.**

This heatmap illustrates the time course proportions of spatial relationships classified as "Same", "Close", "Intermediate" and "Away" between two individuals every 2 hours within groups of four, eight, and 15/16 male and female mice. Hypothetical mouse ID #X and #Y are designated, with the two individuals represented as X-Y to facilitate analysis. The total number of pair combinations is calculated using the mathematical formula ${}_{x}C_{2}$, resulting in 6 combinations for four males and females, 28 for eight males and females, 105 for a group of 15 males, and 120 for a group of 16 females.

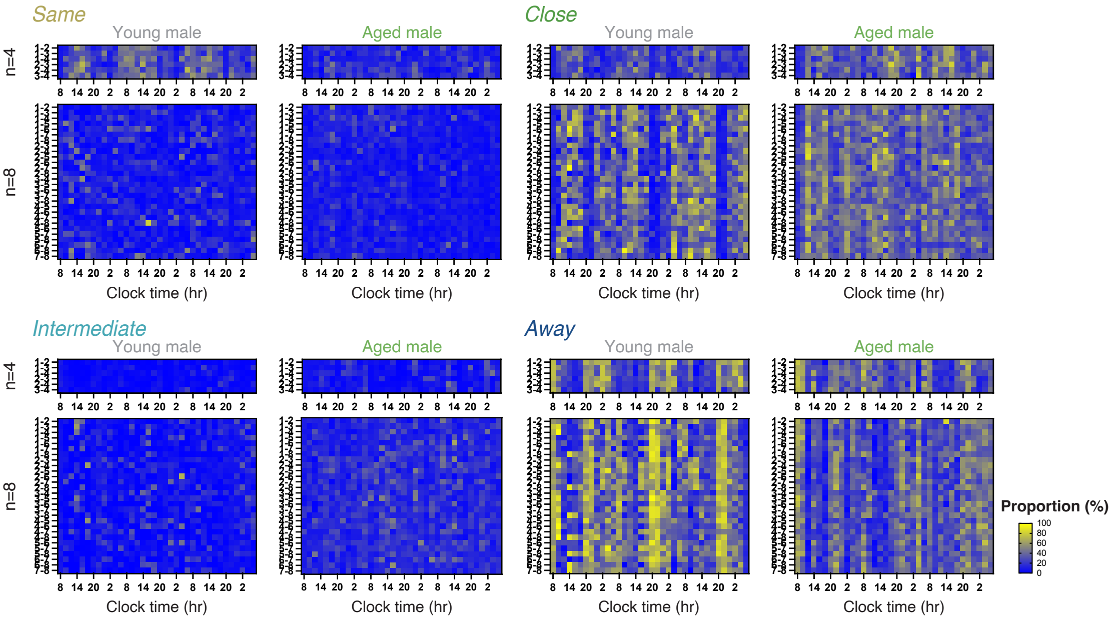


**Supplementary Fig. 20: Analysis of pairwise spatial relationships in young and aged male individuals using the IntelliProfiller workflow.**

This heatmap illustrates the time course proportions of spatial relationships classified as "Same", "Close", "Intermediate" and "Away" between two individuals every 2 hours within groups of four and eight young and aged male mice. Hypothetical mouse ID #X and #Y are designated, with the two individuals represented as X-Y to facilitate analysis. The total number of pair combinations is calculated using the mathematical formula ${}_{x}C_{2}$, resulting in 6 combinations for four young and aged males, 28 for eight young and aged males.

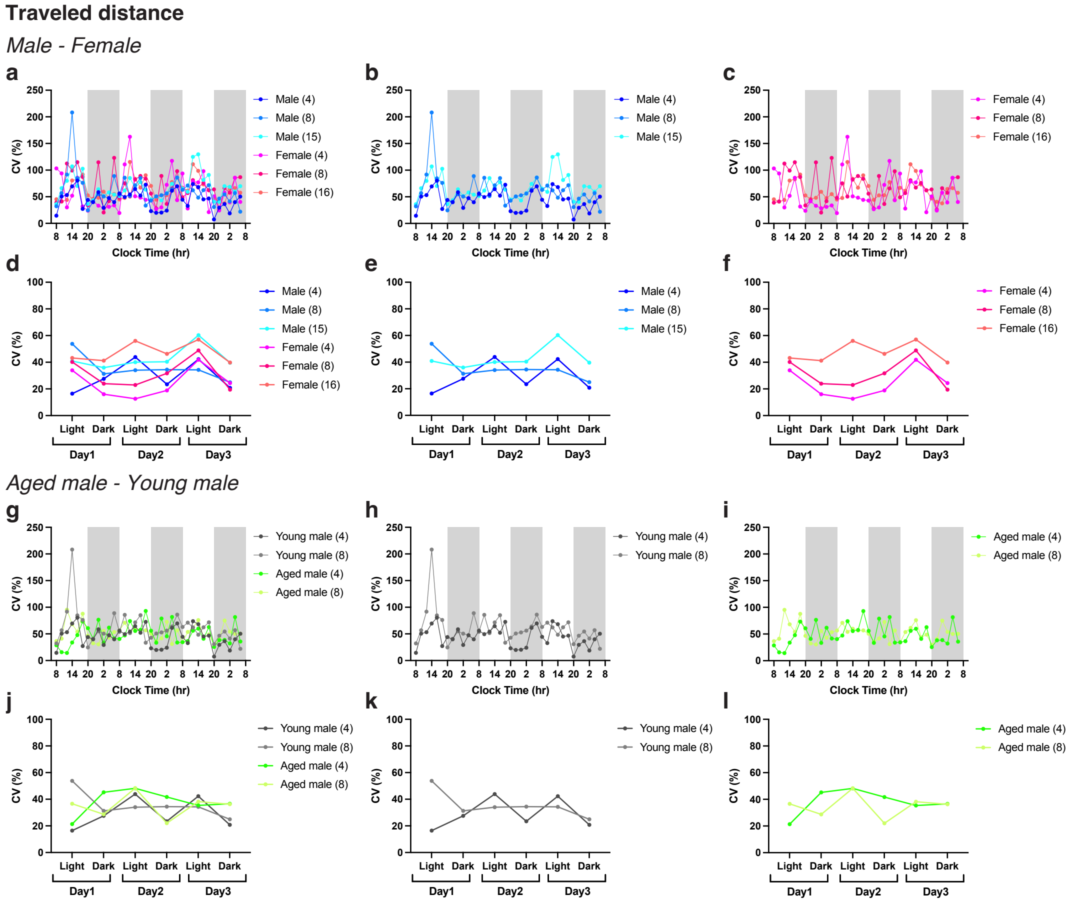


**Supplementary Fig. 21: Coefficient variation (CV) of travel distance in males and females, young and aged males.**

**a-f** CV (%) of travel distance in groups of four, eight and 15/16 males and females (**a, d**), males (**b, e**), females (**c, f**) in 2-hour intervals (**a-c**) and 12-hour intervals (**d-f**). **g-l** CV (%) of travel distance in groups of four and eight young males and aged males (**g, j**), young males (**h, k**), aged males (**i, l**) in 2-hour intervals (**g-i**) and 12-hour intervals (**j-l**).

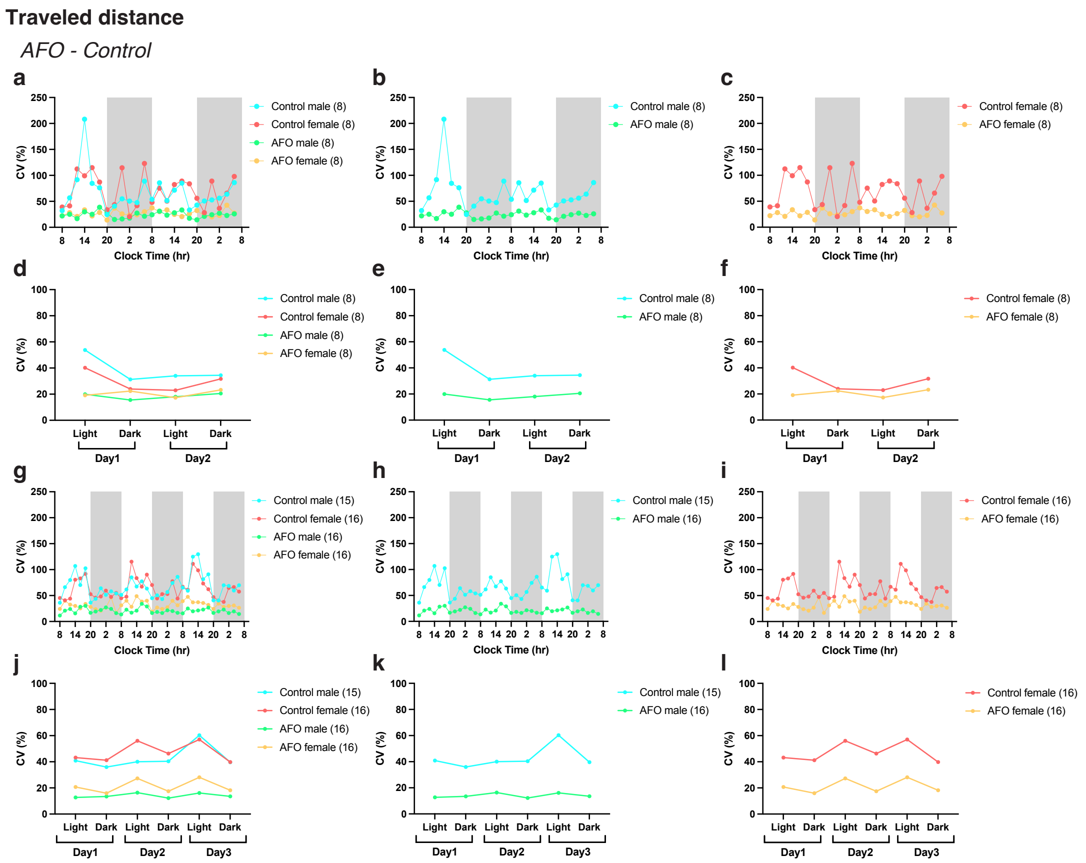


**Supplementary Fig. 22: CV of travel distance in AFO and control males and females.**

**a-f** CV (%) of travel distance in groups of eight AFO and control males and females (**a, d**), eight AFO and control males (**b, e**), eight AFO and control females (**c, f**) in 2-hour intervals (**a-c**) and 12-hour intervals (**d-f**). **g-l** CV (%) of travel distance in groups of 15/16 AFO and control males and females (**g, j**), 15/16 AFO and control males (**h, k**), 16 AFO and control females (**i, l**) in 2-hour intervals (**g-i**) and 12-hour intervals (**j-l**).

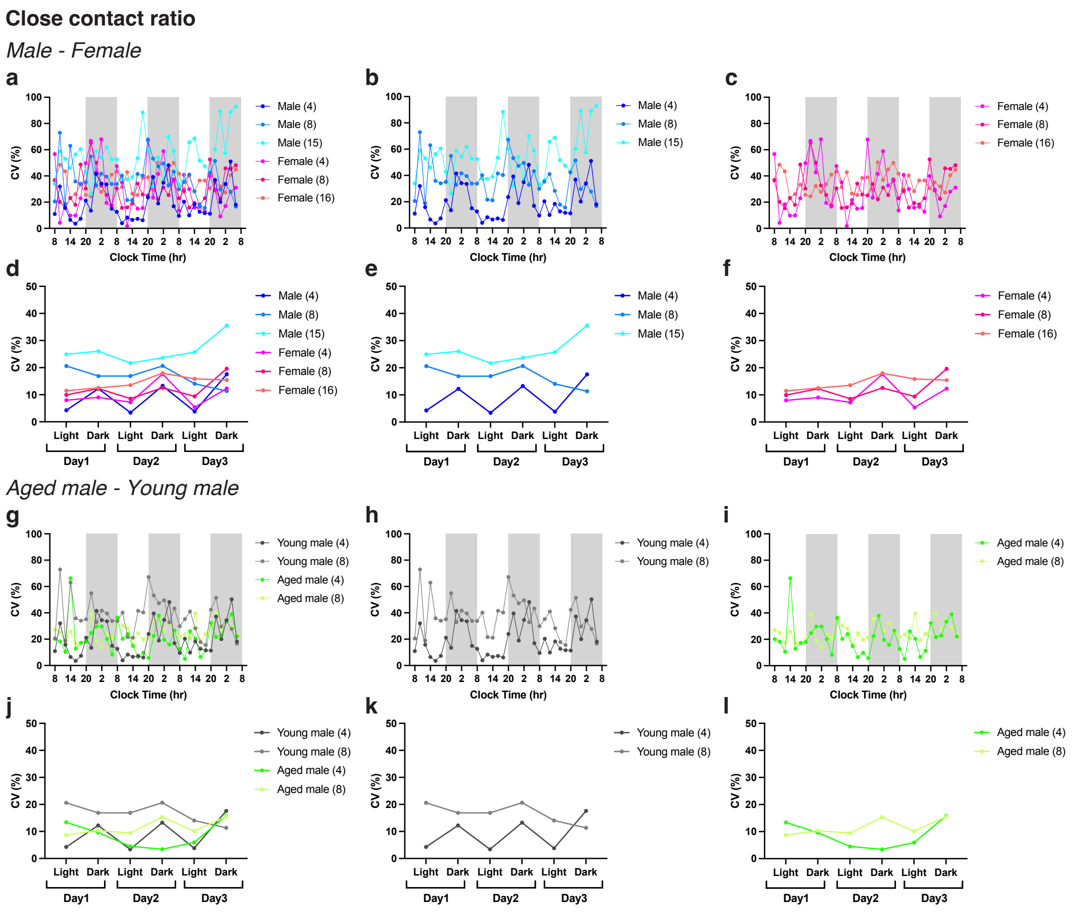


**Supplementary Fig. 23: CV of CCR in males and females, young and aged males.** **a-f** CV (%) of CCR in groups of four, eight and 15/16 males and females (**a, d**), males (**b, e**), females (**c, f**) in 2-hour intervals (**a-c**) and 12-hour intervals (**d-f**). **g-l** CV (%) of CCR in groups of four and eight young males and aged males (**g, j**), young males (**h, k**), aged males (**i, l**) in 2-hour intervals (**g-i**) and 12-hour intervals (**j-l**).

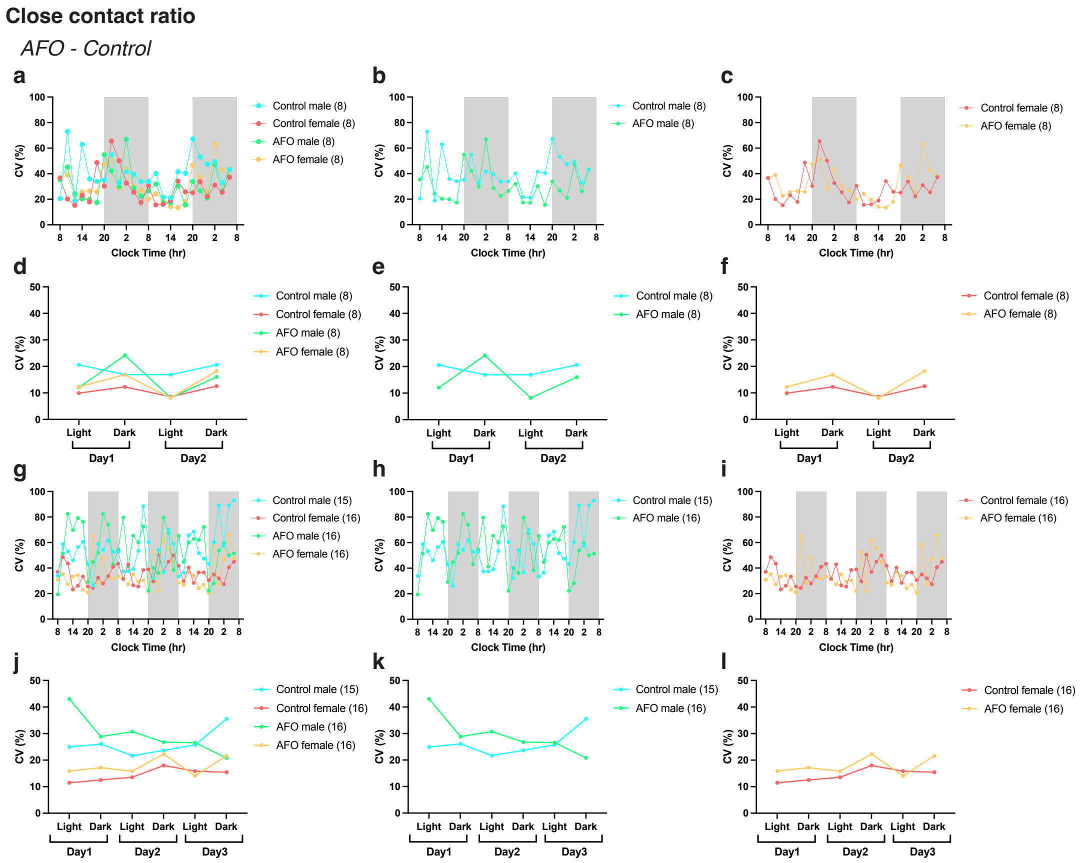


**Supplementary Fig. 24: CV of CCR in AFO and control males and females.**
**a-f** CV (%) of CCR in groups of eight AFO and control males and females (**a, d**), eight AFO and control males (**b, e**), eight AFO and control females (**c, f**) in 2-hour intervals (**a-c**) and 12-hour intervals (**d-f**). **g-l** CV (%) of CCR in groups of 15/16 AFO and control males and females (**g, j**), 15/16 AFO and control males (**h, k**), 16 AFO and control females (**i, l**) in 2-hour intervals (**g-i**) and 12-hour intervals (**j-l**).

**Supplementary Table 1: R and Python packages required for the scripts.**

**a** R packages required for the scripts.

| | Package name | Version | Functionality |
|---|---|---|---|
| 1 | tidyverse | 1.3.1 | Data manipulation, visualization, string operations, data import/export. |
| 2 | openxlsx | 4.2.5 | Reading and writing Excel files |
| 3 | lubridate | 1.7.10 | Manipulation of date and time data |

**b** Python packages required for the scripts.

| | Package name | Version | Functionality |
|---|---|---|---|
| 1 | pandas | 2.2.2 | Data manipulation and analysis |
| 2 | matplotlib | 3.8.4 | Plotting and visualization |
| 3 | scikit-learn | 1.4.2 | Machine learning algorithms and tools (PCA) |
| 4 | numpy | 1.26.4 | Numerical computations |
| 5 | os | built-in | OS interaction and file management |
